# Functional characterization of fatty acid synthase in *Hermetia illucens*

**DOI:** 10.1101/2024.03.22.586266

**Authors:** Yuguo Jiang, Wei Xue, Zhidan Peng, Peng Hou, Zongqing Kou, Bihui Chen, Huimeng Lu, Yongping Huang

## Abstract

Fatty acid synthase (FAS) is a key role in *de novo* lipogenesis (DNL) in most lives. In insects, type I FAS functions in homodimer, activated by a phosphopantetheine transferase (PPT), to *de novo* produce essential saturated fatty acid for cellular processes. In a nova insect species *Hermetia illucens* (Diptera: Stratiomyidae), different from most animals and insects, lauric acid (LA) accumulates to a noticeable level, near content of that in coconut oil or palm kernel oil. However, the origin of LA, from *de novo* biosynthesis or as degraded products from long chain fatty acid, was still ambiguous. Here, in this study, we mined a *FAS* genes (*HiOGS16234*) from *H. illucens* genome and initially characterized its abilities of LA production in a free-fatty-acid producible yeast. After clear the yeast metabolites background of fatty acid by deletion of *ScFAS1* and *ScELO1* sequentially, which responsible for fatty acid genesis and elongation respectively, functions of HiOGS16234 from *H. illucens* was *in vitro* elucidated by analyzing its product portfolio. Through protein domain swap in HiOGS16234, we found the ketoacyl synthase domain and thioesterases domain were responsible for the different product portfolio of HiOGS16234, compared with DmFASN1. Through *in vitro* assay in yeast, the origin of LA in BSF was firstly elucidated at molecular level, which support DNL origin hypothesis in BSF.

## Introduction

As a conserved primary cellular function, *de novo* lipogenesis (DNL) is essential in most lives. Excess acetyl-CoA from carbohydrates took from environment will incorporated into fatty acids or CoA derivatives for further storage in triglycerides or as substrates for membrane components construction (Strable and Ntambi, 2010). Key enzymes in this fatty acid synthesis process are acetyl-CoA carboxylase (ACC) and fatty acid synthase (FAS). ACC catalyzes the limiting step reaction, turning acetyl-CoA into malonyl-CoA, which donates two carbon units in each sequential cycle of condensation in chain elongation (Færgeman and Knudsen, 1997). FASs can be divided into two systems based on architectures. Type I FASs occurs in multifunctional forms in eukaryotes and particular Corynebacterineae, while bacteria and plants synthesize fatty acids with separate, monofunctional enzymes, termed as type II (Leibundgut et al., 2008). Beyond distinct architectures, the process of biosynthesis is highly conserved. Fatty acid synthesis is initiated by transferring acetyl moiety from acetyl-CoA to acyl carrier protein domain (ACP) after activated by phosphopantetheine transferase (PPT), which attaches the prosthetic phosphopantetheine group to ACP (Heil et al., 2019). Then through a repetitive reaction sequence, including condensation, reduction, dehydration, and reduction again, a saturated aliphatic chain with desired length was obtained (Meier and Burkart, 2009). In type I FASs, among domains with distinct enzyme activities, ketoacyl synthase domain (KS), acyltransferase domain (AT) and thioesterases domain (TE) are the key role in chain length control. KS limits the space for chain growing while AT or TE has affinity preference to hydrophobic tail with specific length for transfer or hydrolysis (Heil et al., 2019; Herbst et al., 2018).

In insects, the ability of DNL was long thought to be ubiquitous as other animals. There were thus little reports about fatty acid biosynthesis or DNL related enzyme activity in last decades and more work was focused on physiological phenotypes in insects, including growth, development, oviposition, and desiccation tolerance (Alabaster et al., 2011; Chung et al., 2014; Li et al., 2016; Moriconi et al., 2019; Parvy et al., 2012; Pei et al., 2019; Song et al., 2022). One hot spot of DNL research in insects was about the ability of lipid synthesis in wasps due to their parasitic lifestyle (Prager et al., 2019; Visser et al., 2017, 2010). In much early years, between 1960s and 1990s, fatty acid synthesis in Dipteran insect *Drosophila melanogaster* was focused to investigate from compositions to metabolism (Church and Robertson, 1966; Kato, 1965; Keith, 1967, 1966; Keith et al., 1967). They found in fruit flies’ neutral lipids, there were small amount of medium chain fatty acid (MCFAs), including lauric acid (LA, C12) and myristic acid (C14) (de Renobales and Blomquist, 1984). Then after purified its fatty acid synthase and did *in vitro* assays, they found that *D. melanogaster* had inherent capability to produce MCFAs, especially for myristic acid (C14) (de Renobales et al., 1986; de Renobales and Blomquist, 1984).

In another Dipteran insect *Hermetia illucens* (Diptera: Stratiomyidae), acted as an organic waste recycler, lauric acid mostly accumulates to a surprising level (36%-60%)(Almeida et al., 2020), which is uncommon in other insects, such as flies, crickets, locusts, mealworms and superworms (Adámková et al., 2017; Cripps et al., 1986; Jayanegara et al., 2020; Suryati et al., 2023; Thompson, 1973; Verheyen et al., 2018). Meanwhile, fed with various common organic substrates with little LA, including chicken feed, manure, vegetables, fruits and cereal, BSF larvae showed a relative constant lipids profiles and relative high LA content (Almeida et al., 2020; Guil−Guerrero et al., 2020; Rodrigues et al., 2022). As a main component of coconut oil or palm kernel oil (Dayrit, 2015) appeared in a terrestrial insect with a relative high content, a DNL origin hypothesis was proposed and tested in different ways. Transcriptomic analysis through BSF larvae development revealed that putative *HiFAS* and *HiACC* were significantly upregulated through the whole larval stage after hatching, especially highly expressed around the third instar stage, which implied that LA may from native biosynthesis in BSF larvae (Zhu et al., 2019). Deuterium labelling assay provided a closer insight into the molecular mechanism of fatty acid biosynthesis in BSF. In two different complex diets with deuterated water, LA was almost entirely found in deuterated form, which indicated LA was from DNL (Hoc et al., 2020). Meanwhile, as core gene of DNL, the putative *HiFAS* and *HiACC* genes were partially characterized in incomplete sequences without any *in vivo* or *ex vivo* functional identification (Giannetto et al., 2020). Until now, only one gene, *HiACP*, which participated in fatty acid biosynthesis was functionally characterized (Peng et al., 2023). The DNL core genes, especially for *HiFAS*, were needed to be directly investigated at molecular level to test the DNL origin hypothesis of LA.

In this study, to characterize the role of *HiFAS*s in LA accumulation, we cloned two *HiFAS* candidates and overexpressed heterologously in yeast for functional verification, by the way, together with two putative Dipteran *PPT*s. In a free-fatty-acid producible yeast, one of the *HiFAS*s (*HiOGS16234*) could bring significant LA after overexpression. Further diminishing the fatty acids produced by yeast itself through metabolic manipulation, we proved that HiOGS16234 indeed produced more LA than DmFASN1 in yeast, which corresponded with the LA content in both flies. By swapping several domains, which may be responsible for chain-length control of fatty acid, we found that the ketoacyl synthase domain and thioesterases domain took the main responsibility for the distinct product portfolio of HiOGS16234.

## Materials and Methods

### Materials and strains

The laboratory-maintained wild-type line of BSF, which was originally collected from Dr. Ziniu Yu’s lab, underwent inbred crossing for generations in our lab (Zhan et al., 2019). *S. cerevisiae* strain (IMX581) purchased from EUROSCARF was used as parent strain for all constructs. *E. coli* strain DH5α was used for gene cloning. *SFP* (UniprotKB accession no. P39135.2) was synthesized by Synbio Technologies (Suzhou, China). All the strains used or constructed in this study are listed in Table S2 and the primers used for the construction are listed in Table S1.

### Clone of Dipteran *FAS* genes and *PPT* genes

Candidate *HiFAS* and *HiPPT* genes were mined from genome of *H. illucens* (Generalovic et al., 2021; Zhan et al., 2019) using *DmFASN*s (*CG3524*, *CG3523*, *CG17374*) and *SFP* as query, respectively. Candidate *DmPPT* gene was also mined by *SFP* in FlyBase (https://flybase.org). Polymerase chain reaction (PCR)-amplification was applied to obtain all sequences from corresponding complementary DNA (cDNA) of larval fat body. For cDNA synthesis, total RNA was isolated from fat body in 3^rd^ larval instar of BSF using TRIeasyTM Total RNA Extraction Reagent (Yeasen, Shanghai, China). One µg of total RNA was processed using PrimeScript RT reagent Kit with gDNA Eraser (Takara, Beijing, China). Amplified products were sequenced and verified after cloning into a pJET1.2-T vector through CloneJET PCR Cloning Kit (Thermo Scientific, Waltham, USA).

### Phylogenetic analysis of *HiFAS*s

Protein sequences of candidate FASs from seven typical Dipteran insects (*Lucilia cuprina*, XP_023298940.2; *Musca domestica*, XP_019894443.2; *Anopheles gambiae*, XP_319941.4; *Culex quinquefasciatus*, EDS45473.1; *Aedes aegypti*, XP_001658180.2; *Aedes albopictus*, XP_029715180.1) were collected from NCBI by BLASTP against Genbank nonredundant protein database (nr) using CG3523 from *D. melanogaster* as a protein query. Together with a Lepidopteran FAS (*Bombyx mandarina*, XP_028030558.1) as reference, all the sequences were aligned using MAFFT, and neighbor-joining tree was constructed using the program MEGAX with JTT matrix as substitution model, pairwise deletion for gaps and 1000-replicate bootstrap analysis.

### Construction of plasmids and yeast strains

For co-expression *FAS*s and *PPT*s, the vector used was pESC-URA (Agilent, USA), containing *GAL1* and *GAL10* yeast promoters in opposing orientation. Dipteran *FAS*s were under control of *GAL1p* and PPTs were under *GAL10p* control. For *in vivo* homologous recombination in yeast, the adjacent fragments made up the vector all shared ∼ 40 bp identical sequences. In domain swap assay, the KS/AT/TE fragments were annotated after alignments (Fig. S4). Transformation of DNA fragments was conducted using LiAc/SS carrier DNA/PEG method (Gietz and Schiestl, 2007). Transformants were selected on selective plates without uracil and confirmed by sequencing.

### Strain cultivation and metabolites extraction

For *URA3* marker selection, yeast clones were grown in synthetic complete (SC) medium composed of 6.7 g/L yeast nitrogen base without amino acid (BD Difco, Sparks, USA) and 20 g/L glucose. For auxotroph strains, 0.5 mM pentadecanoic acid (C15) or hexadecanoic acid (C16) was dispersed in the medium by nonionic surfactants Brij58 or NP40 (0.5% w/v) after sterilization by filtration.

Shake flasks fermentation was conducted in triplicates in synthetic medium consisting of 2 g/L (NH_4_)_2_SO_4_, 12 g/L KH_2_PO_4_, 0.5 g/L MgSO_4_·7H_2_O, 20 g/L glucose, 0.1% (v/v) trace mental solution and 0.1% (v/v) vitamin solution (Zhou et al., 2016). Inoculated at an initial OD_600_ of 0.1 in 10 mL synthetic medium, culture was cultivated at 250 rpm, 30 °C for 72 h (for strain YG1 series) or 96 h (for *Scfas1*Δ or *Scfas1 Scelo1*Δ strains).

For free fatty acids extraction from strain YG1 series, simultaneous extraction and methylation was conducted as described previously (Jiang et al., 2022). Briefly, cell cultures from flask fermentation were diluted by 2-fold. Then, (C_4_H_9_)_4_N^+^OH^-^ was added into vials followed with CH_2_Cl_2_ solution containing CH_3_I and heptadecanoic acid as an internal standard. After extraction, organic phases were evaporated to dryness and suspended in hexane for further analysis.

For total fatty acid extraction from *Scfas1*Δ or *Scfas1 Scelo1*Δ strains, cell pellets were collected by centrifuging 10 mL cell culture from flask fermentation. After washed by ddH_2_O and vacuum freezing drying for overnight, cell pellets were weighted and transferred into 3 mL mixture of 10% BF_3_-MeOH and hexane in ratio of 2:1 (v/v) for methylation in 80℃ for 4h. The organic phase was dehydrated by anhydrous Na_2_SO_4_ for further analysis.

### BSF rearing, raw fat extraction and fatty acids methylation

For laboratory diet, the newly hatched BSF larvae were reared in moistened wheat bran (1:2 w/v of dry matter/ddH2O) under an 8:16 L:D photoperiod at 28℃ with relative humidity of 50%-60%. The feeding materials were refreshed every 24-48 hours until over 75% of larvae reached prepupal stage. The larvae collected at 7 days post hatching were defined as L4 instar, which were separated from the remaining residues and dried in 75℃ for overnight before raw fat extraction.

For kitchen waste diet, 0.8 g newly hatched BSF larvae were incubated in 2.5 kg kitchen waste collected from the canteen, mixed with 1 kg wheat bran to adjust the moisture for larvae growth. One or a half kilo kitchen waste were added every 24-48 hours, based on the bioassimilation progress, until over 75% of larvae reached prepupal stage. The larvae collected at 6 days, 11 days and 15 days post hatching were defined as L4, L5 and prepupae, respectively. The larvae also were separated from the residues and dried in 75℃ until constant weight before raw fat extraction.

The raw fat extraction of BSF larvae was determined by the standard protocol. Briefly, 1 g of the dried sample, from laboratory or kitchen waste diet, was ground and subjected to Soxhlet extraction with n-hexane at 75℃ overnight. After vacuum evaporation, 50 mg extracts were transferred into 1 mL of NaOH-MeOH solution (2%, w/v) for saponification. Then, 1mL 10% BF_3_-MeOH was added for methylation, followed with 2 mL hexane for extraction. The fatty acid methyl esters were determined by gas chromatography.

### Chemical analysis of metabolites

For fatty acid profiles of BSF larvae, an GC2010 pro (Shimazu, Suzhou, China) was equipped with a TG-WaxMS capillary column (60 m × 0.25 mm × 0.25 µm, Thermo scientific) and a flame ionization detector (FID). The GC program was as follows: initial temperature of 120 °C; ramp to 240℃ at rate of 5℃/min, maintain 6 min. The temperature of inlet and detection were 220℃ and 240℃, respectively. The sample volume of per injection was 2 µL in split ratio of 40. The flow rate of carrier gas (nitrogen) was kept at 1.5 mL/min.

For fatty acid profiles of yeast strain, a TG-5MS capillary column (30m × 0.25 mm × 0.25 µm, Thermo Scientific) was equipped. The GC program was as follows: initial temperature of 120°C; ramp to 165°C at rate of 30°C /min; then ramp to 212°C at rate of 2°C /min. The temperature of inlet and detection were 250°C and 260°C, respectively. The sample volume of per injection was 2 µL in split ratio of 20. The flow rate of carrier gas (nitrogen) was kept at 1.25 mL/min.

Qualitative analysis was conducted by authentic compounds purchased from Sigma. Fatty acid relative content was quantified by percentage of mol of each fatty acid in extracts, which was calculated by external standard method.

### Statistical analysis

A one-way analysis of variance (ANOVA) with Tukey’s multiple comparisons test was used to analyze the differences between results unless otherwise noted. Bar charts were drawn and analyzed with GraphPad PRISM 8.0 software (San Diego, California, USA). Three independent biological replicates were used for each test. Error bars represent the standard deviation (S.D.).

## Results

### 1. Characterization of *HiFAS* candidate genes in BSF genome

In *H. illucens*, larvae fed in food waste could accumulate LA more than 40% in the grease at prepupae stage (Fig. 1A). Combined with relative constant fatty acid profiles in different feeding assays (Rodrigues et al., 2022), we hypothesized that larvae of *H. illucens* could produce LA by *de novo* lipogenesis. The laboratory rearing material, wheat bran, was firstly detected to rule out the possibility of LA introduction from the fed. After 12 days fed by wheat bran, the larvae accumulated substantial LA and little myristic acid (C14, MA) while there was no LA or MA detected in wheat bran oil extracts (Fig. 1B), which means at least these MCFAs were not directly provided by the feeding materials.

**Fig 1.**
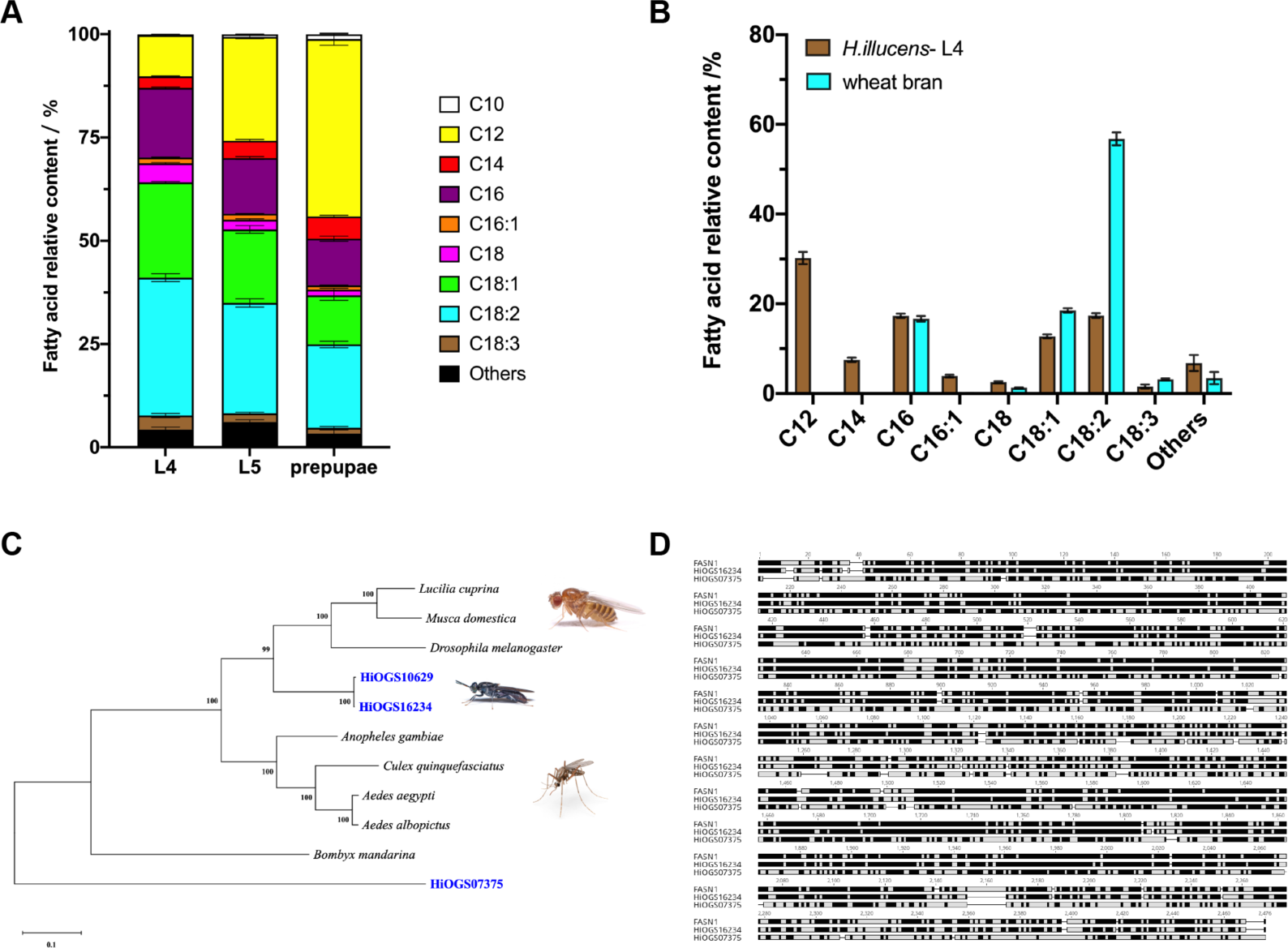
Cloning of *HiFAS* candidate genes from BSF genome. (A) Fatty acid profiles in different larval stage of BSF fed with kitchen wastes. (B) Fatty acid profiles of BSF larvae in the 4^th^ larval instar fed with moisture wheat bran, compared with that of the feeding materials. (C) Phylogenetic analysis of FAS protein sequences from BSF and other typical Dipteran insects. FAS candidates from BSF are shown in blue. (D) Alignment analysis in protein sequences of two HiFAS candidates and DmFASN1.

In *D. melanogaster*, FASN1 (CG3523) has been functionally verified to mainly produce long-chain fatty acids (LCFAs) with a small amount of MCFAs by substrate incorporation with purified FAS fractions (de Renobales and Blomquist, 1984). Taking *DmFASN1* as a query sequence, we mined three *HiFAS* candidate genes in annotated genome (*HiOGS16234*, *HiOGS10629* and *HiOGS07375*). Among them, HiOGS10629 show a high protein sequence identity with HiOGS16234 (95.97%, Fig. S1). The protein sequence similarity between each of them with DmFASN1 were 89.44%, 85.91% and 71.29%, respectively. In phylogeny analysis, HiOGS16234 and HiOGS10629 were located between cluster of flies and mosquitos (Fig. 1C), which was well corresponded with whole genome phylogenetic relationship. However, HiOGS07375 located even further than reference sequence from moth (Fig. 1C), which implied that the enzyme function of HiOGS07375 may not be similar with DmFASN1. After several cloning trials, whole length of *HiOGS16234* and *HiOGS07375* was cloned from cDNA library of L3 fat body. The protein length of both HiFAS candidates were 2416 aa and 2386 aa, respectively, which was near the length of DmFASN1(2438 aa) (Fig. 1D). To simplify illustration, the names of both candidates were abbreviated into HiOGS1 and HiOGS0, respectively.

### 2. Overexpression of *HiFAS* candidate genes in FFA producible yeast strain

To verify the enzyme activity of *HiFAS* candidates, we designed an *in vitro* overexpression assay (Fig. 2A). Budding yeast (*Saccharomyces cerevisiae*), produces little MCFA natively, was used as chassis to load with a gene cassette on a vector including a *FAS* and a phosphopantetheinyl transferase gene (*PPT*). After fermentation in minimal medium, fatty acid profiles were determined by metabolites analysis. In the cassettes, as an essential cofactor, PPT mediates the transfer of pantetheine arm form CoA to the acyl carrier protein domain (ACP) of animal FAS to activate it into *holo*-synthases (Beld et al., 2014). If PPT cooperate well with specific FAS, the featured products beyond yeast metabolic background will appear——for DmFASN1, which is myristic acid (C14) and LA for HiFAS candidates. However, in *H. illucens*, the *PPT* has not been characterized and putative *PPT* in *D. melanogaster* (*Ebony*) was not proved to act in DNL(Richardt et al., 2003). We firstly used *SFP*, an extensively implemented *PPT* from *Bacillus subtilis* (Leber and Da Silva, 2014; Mootz et al., 2001), as query sequence to obtain native *PPT* candidates in both Dipteran insects. Then, as catalytical function of *DmFASN1* has been well characterized as reported, we co-expressed *SFP* or *DmPPT* with *DmFASN1* to evaluate the effectiveness of this *in vitro* overexpression assay in a free fatty acid producible strain YG1. Compared with empty vector, both co-expression groups produced substantial myristic acid and small amount of LA (Fig. 2B).

**Fig 2.**
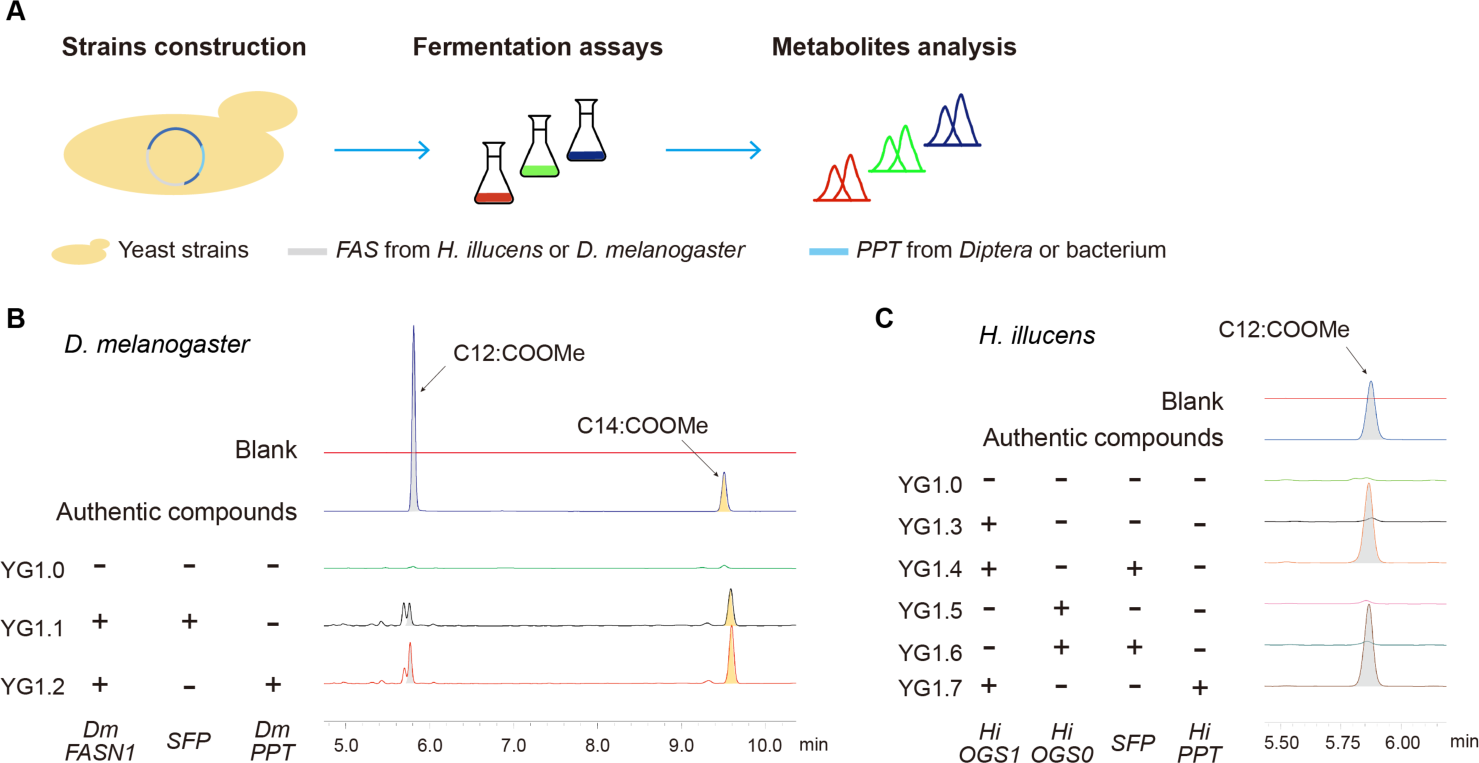
Overexpression of Dipteran *FAS* genes in FFA producible yeast strain. (A) Workflow of the *in vitro* overexpression assay to characterize Dipteran *FAS*s in yeast. (B) Gas chromatography of free fatty acid producible strains loaded with *DmFASN1*. (C) Gas chromatography of free fatty acid producible strains loaded with *HiFAS*s.

*HiOGS1* and *HiOGS0* were also co-expressed with *SFP* firstly. Within our expectation, *HiOGS1*, whose protein sequence located much nearer from DmFASN1 than HiOGS0 in phylogenetic relationship analysis, produced amounts of LA, while nearly nothing around the same retention time in *HiOGS0* group, as similar as the control group (Fig. 2C). The *HiPPT*, mined from the BSF genome, also was tested under the same condition. The putative PPT from BSF could also activate HiOGS1 to produce comparable LA as SFP did (Fig. 2C). Combined with transcriptome results from BSF, the products from the *in vitro* overexpression assay suggested that BSF was able to produce LA through *de novo* lipogenesis, which partially elucidated the origin of LA in BSF.

### 3. Functional replacement of ScFAS1 by Dipteran FASs

In budding yeast, besides taking from medium, fatty acyl compounds also could be *de novo* synthesized from acetyl CoA by native ScFAS1/2 (Schüller et al., 1992). Different from animal FASs, fungal FASs work in a hetero-hexamer, comprising six α- and six β-chains, encoded by *ScFAS1* and *ScFAS2*, respectively (Heil et al., 2019). Given that in above *in vitro* overexpression assay, fatty acids were produced by native ScFASs and Dipteran FASs, *ScFAS1* or *ScFAS2* need to be deleted to block the FA synthesis by ScFASs for a clear metabolic background. In strain YG3, *ScFAS1* was deleted to abolish the native *de novo* lipogenesis based on YG2. This strain reserved the ability to take lipids from medium, so it survived as noticeable dots on YPD medium (Fig. 3A). When even odd-chain fatty acids, which rarely synthesized by yeast, were added into the medium, dispersed by NP40, the growth defect caused by *ScFAS1* deletion was nearly fully rescued (Fig. 3A). In liquid medium, YG3 also could not accumulate enough biomass without any extra FAs addition (Fig. 3B).

**Fig 3.**
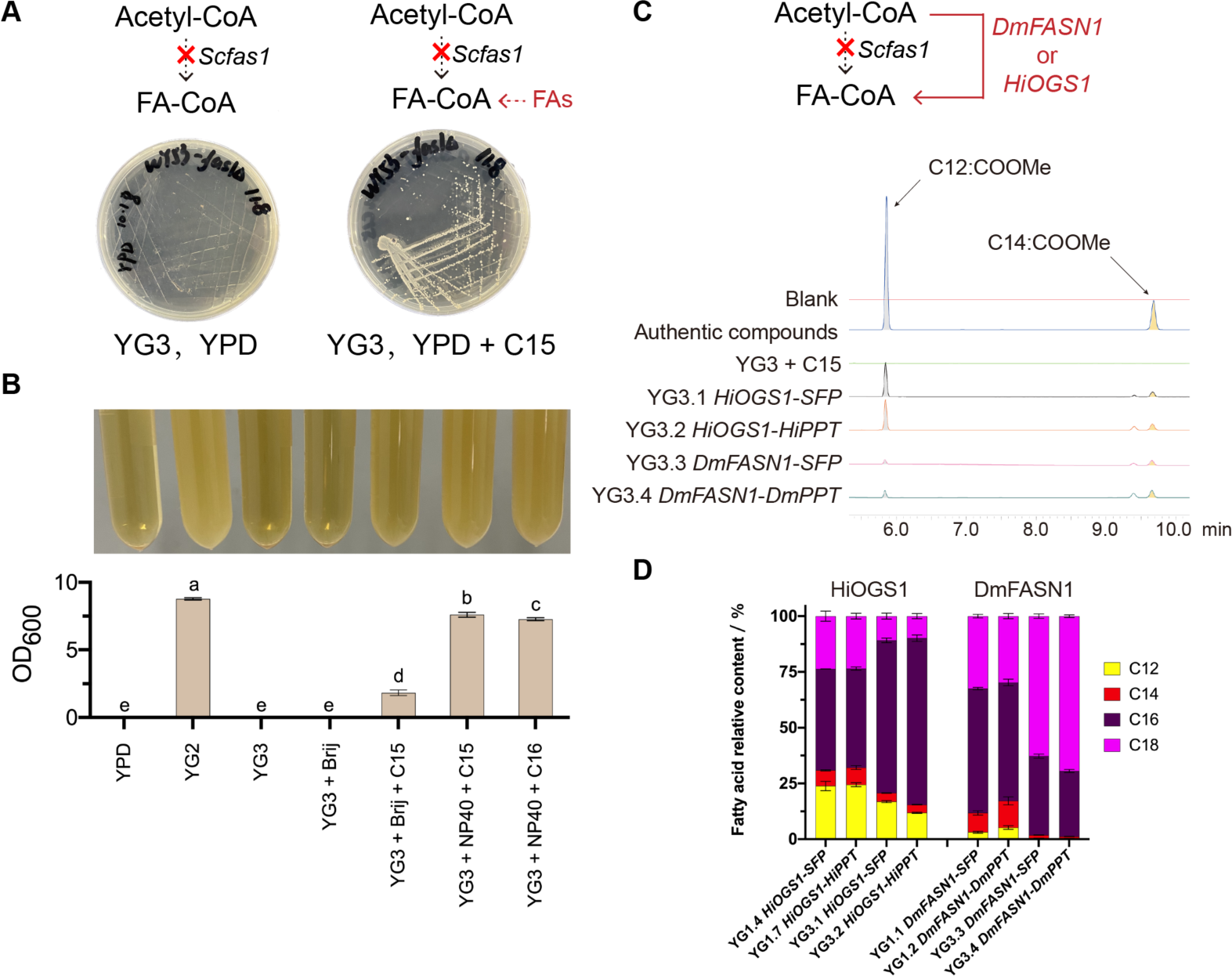
Functional replacement of ScFAS1 by Dipteran FASs. (A) Construction of *Scfas1*Δ strain. (B) Rescue of growth defect in *Scfas1*Δ strain by pentadecanoic acid (C15) addition in liquid medium. (C) Gas chromatography of *Scfas1*Δ strains survived by pentadecanoic acid (C15) addition or Dipteran *FAS*s cassettes overexpression. (D) Changes in fatty acid relative content of strains loaded with Dipteran *FAS*s cassettes before and after *ScFAS1* deletion. In B and D, the standard deviation was calculated from biological replicates. In B, different letters indicate statistically significant differences across groups using a one-way analysis of variance (ANOVA) with Tukey’s multiple comparisons test, *P* < 0.05

In strain YG3, we further introduced the Dipteran *FAS* overexpression cassettes to rescue the growth defect by endowing the strain with lipogenesis ability. Without any extra FAs addition, the resulted strains accumulated comparable biomass as FAs added group (Fig. S2A) and produced featured compounds (Fig. 3C). Compared with corresponding YG1 strains, however, the YG3 strains with gene cassettes had much higher proportions of LCFAs (Fig. 3D), which implied that the native fatty acyl elongation had changed the product portfolio of heterologous FAS under growth stress brought by *Scfas1*Δ.

### 4. Verification of Dipteran FASs product portfolio by blocking elongation

In budding yeast, there are three fatty acyl elongases, ScELO1/2/3. Among them, ScELO1 extends medium-chain fatty acids to long-chain fatty acids by incorporation of malonyl-CoA, while ScELO2/3 are involved in sphingolipid biosynthesis and act on fatty acids of up to 24 carbons in length (Dittrich et al., 1998; Oh et al., 1997; Toke and Martin, 1996). To block the native fatty acyl elongation of potential MCFAs produced by Dipteran FASs, *ScELO1* was deleted in strain YG3 to form strain YG4. Given both strains only grew in minimal medium with extra FAs added, 0.5 mM pentadecanoic acid (C15) was dispersed in the medium to rescue this nutrition deficiency. Compared with strain YG3, further impaired fatty acyl modification caused severer growth defects in strain YG4, either after FAs addition or *FAS* cassettes overexpression (Fig. S2B). Despite poor growth, without any extra FAs added into the minimal medium, strain YG4 loaded with Dipteran *FAS*s survived and as expected, less pentadecanoic acid was elongated into heptadecanoic acid after *ScELO1* deletion, meanwhile more pentadecenoic acid (Z10-15:COOH) was produced (Fig. S3), which corresponded with reported before.

Little native fatty acyl elongation in strain YG4 clear the mist over the product portfolio of both Dipteran FASs. Strain YG4.1 and YG4.2, loaded with DmFASN1, produced considerable myristic acid (C14) while Strain YG4.3 and 4.3, loaded with HiOGS1, synthesized much more lauric acid (C12) and myristic acid (C14) (Fig. 4BD), compared with strains possessing complete elongation pathway (Fig. 4CE). In wild type strain, long-chain fatty acids, such as palmitic acid (C16), stearic acid (C18) and their corresponding unsaturated acids, account for over 80% of all acids (Blagovi et al., 2001). Although heterologous FASs or extra FAs addition could partially rescue the auxotroph strains, quite distinct fatty acid profiles put the result strains under much stress, to the extent that results strains accumulated significantly little biomass, especially for strains loaded with HiOGS1 cassettes, whose fatty acid profiles were more altered (Fig. S2B).

**Fig 4.**
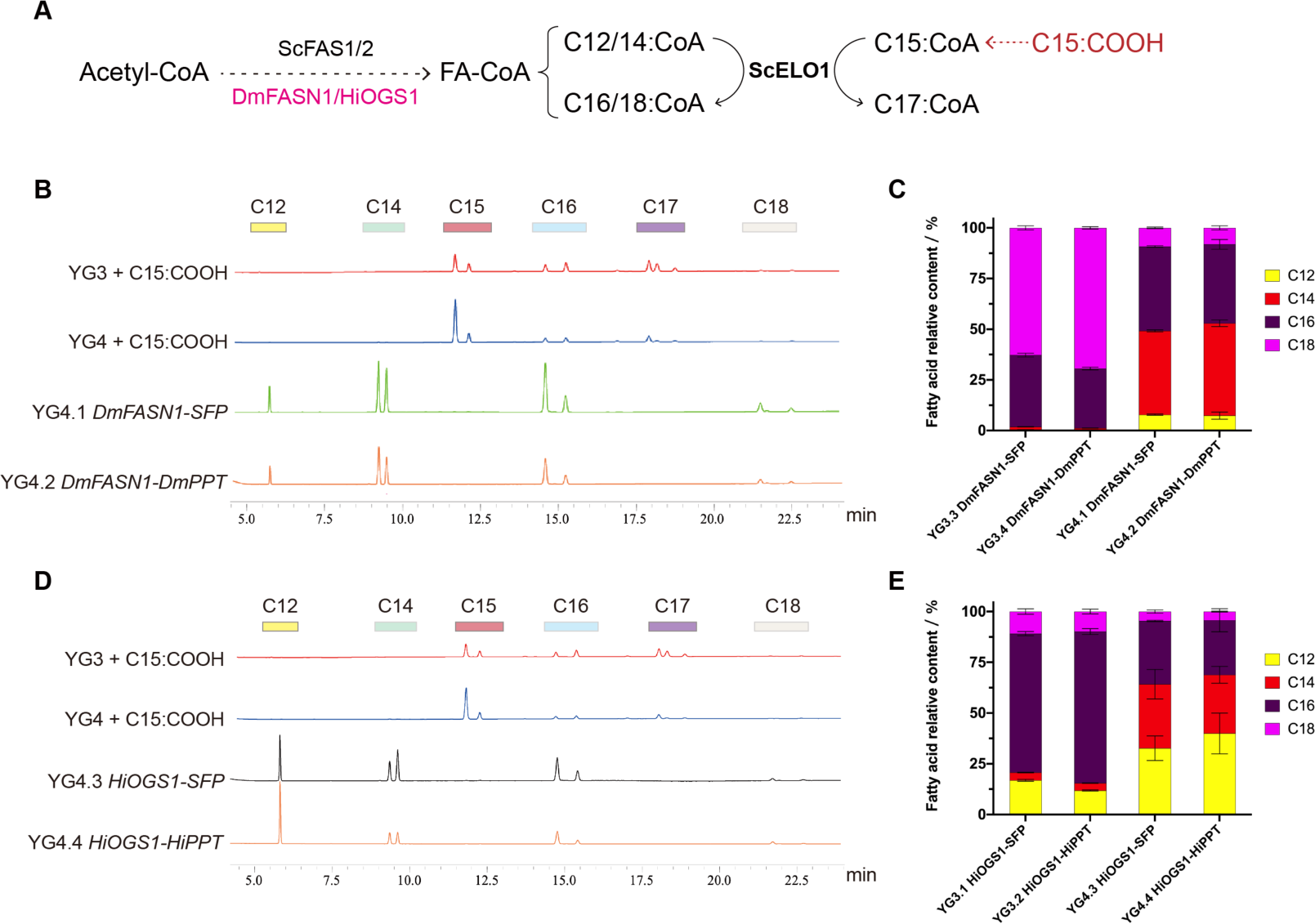
Blockage in native fatty acyl elongation pathway. (A) Scheme for native fatty acyl elongation pathway in yeast. (B) Gas chromatography of *Scelo1*Δ and *Scfas1*Δ strains rescued by pentadecanoic acid (C15) addition or *DmFASN1*(B) and *HiOGS1*(D) cassettes overexpression. (C) Changes in fatty acid relative content of strains loaded with *DmFASN1*(C) or *HiOGS1*(E) cassettes before and after *ScELO1* deletion. In C and E, the standard deviation was calculated from biological replicates.

### 5. Domain swap for chain-length regulation parts

Although HiOGS1 shared 89.44% similarity and 70.88% identity of protein sequence with DmFASN1, the fatty acid profiles of HiOGS1 revealed by *in vitro* assay were distinct from that of DmFASN1. In strain YG4.3 or YG4.4, either HiPPT or SFP as cofactor, HiOGS1 produced nearly equal proportions of lauric acid (C12), myristic acid (C14) and palmitic acid (C16) (Fig. 5A), around 30%. By contrast, major products of DmFASN1 were myristic acid (C14) and palmitic acid (C16), accounted for 43% and 40% on average, respectively (Fig. 5A). The tendency to produce MCFAs, especially for lauric acid, may attributed by specific domains in HiOGS1.

**Fig 5.**
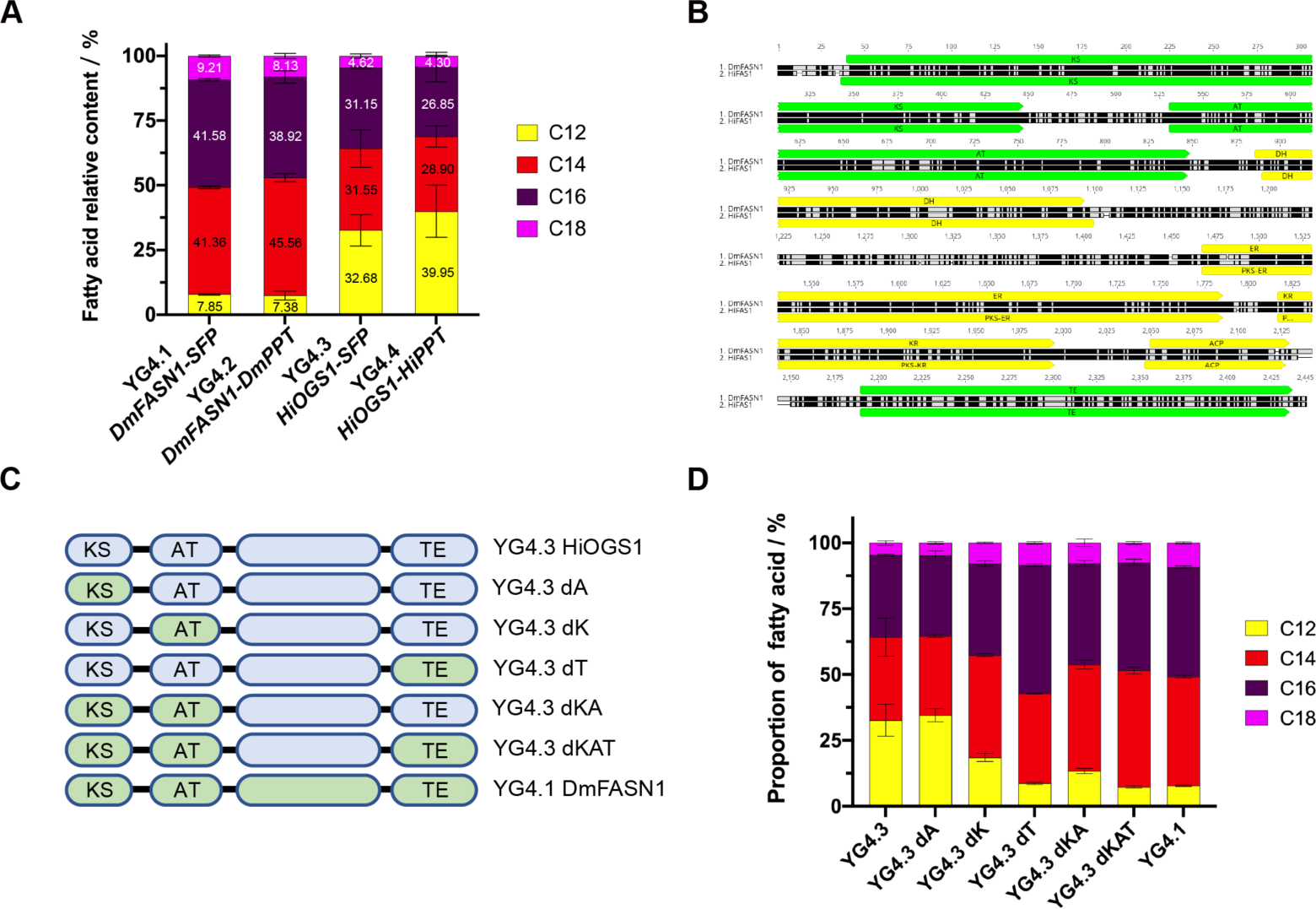
Domain swap in HiOGS1. (A) Comparison of fatty acid relative content of strains loaded with different Dipteran *FAS* cassettes. (B) Alignments of HiOGS1 and DmFASN1 with domain annotations. Potential chain-length regulation domains are shown in green and other domains are in yellow. (C) Design of domain swap in HiOGS1. (D) Changes in fatty acid relative content of strains with chimera *HiOGS1*. In A and D, the standard deviation was calculated from biological replicates.

We focused on chain-length regulation domain: ketoacyl synthase domain (KS), acyltransferase domain (AT) and thioesterase domain (TE) (Fig. 5B). We hypothesized that these domain in HiOGS1 shortened the chain-length of fatty acids into MCFAs, especially for lauric acid, so we designed a domain swap in HiOGS1, in which corresponding domains in DmFASN1 were swapped into HiOGS1 to detect its catalytical functions (Fig. 5C). By single domain swap, we found that AT domain from DmFASN1 did not bring any change in fatty acid profiles in HiOGS1, while KS and TE domain swap significantly decreased the content of lauric acid (Fig. 5D and S5A) and increased the weighted chain length of fatty acids in both strains (Fig. S5B). When all these domains from DmFASN1 were swapped into HiOGS1, strain YG4.3 dKAT showed a much similar fatty acid profiles as strain with DmFASN1 (Fig. 5D and S5A). and the weighted chain length of strain YG4.3 dKAT inevitably showed no significance compared with strain YG4.1 (Fig. S5B). At a holistic viewpoint, the replacement of KS or TE domain in HiOGS1 turned the fatty acid profiles into a DmFASN1-like pattern ((Fig. S5C), which suggested that in HiOGS1, KS and TE domain are the chain-regulation parts, which partially elucidated molecular mechanisms of the origin of lauric acid in BSF.

## Discussion

As reported, partial sequences of *HiFAS* genes had been characterized by transcriptomic analysis or quantitative gene expression (Giannetto et al., 2020; Zhu et al., 2019). Verified by their primers provided in supplementary materials, one of the mentioned putative *FAS* was corresponded with *HiOGS16234* in this study (Zhu et al., 2019) and the other was *HiOGS07375* (Giannetto et al., 2020). In this study, we obtained the whole sequences of two putative *HiFAS* genes and completed functional characterization of them, together with two Dipteran *PPTs*. By an *in vitro* overexpression assay, we found that *HiOGS16234* could produce amounts of LA in yeast and after further metabolic engineering, the product portfolio was unfolded (Fig. 4DE), which supported hypothesis about DNL origin of LA in BSF. By domain swap in HiOGS16234, we further created a chimera HiFAS (dKAT), which showed high similarity to DmFASN1 in fatty acid profiles (Fig. 5D and Fig. S5C). Domain swap assays revealed that KS and TE domain were the key role in chain length regulation in HiFAS, compared with DmFASN1.

In *D. melanogaster*, DmFASN1 was functionally verified the catalytic activity with *in vivo* incorporation of labelled acetate and *in vitro* incorporation of labelled acetyl-CoA and label free malonyl-CoA in ratios. Fatty acid profiles of chain length distribution in *in vivo* assay was similar to that in yeast (Fig. 5A) (de Renobales and Blomquist, 1984). However, in *in intro* assays, the fatty acid profiles were dramatically changed by artificially set of ratio between acetyl-CoA and malonyl-CoA and ionic strength, comparable with *in vivo* results only under specific conditions (de Renobales et al., 1986; de Renobales and Blomquist, 1984), which meant to some extent, product portfolio of FAS was determined by specific outer micro environment surrounding the enzyme. For HiFAS, there was even no such results from *in vivo* assays could performed as references. Thus, the products portfolio of HiFAS was also a reflection by yeast, which may be distorted by designed *in vitro* assays. The real catalytical activity of HiFAS need more evidence to support.

As domain swap revealed the importance of KS and TE domain in HiFAS, we still could not explain the difference between the fatty acid profiles of DmFASN1 and HiFAS in details. Since the chimera HiFAS (dKAT) showed a similar profiles as DmFASN1, further structure analysis could be conducted to elucidate the molecular mechanisms at amino acid level. Furthermore, as *FAS* gene conserved among insects, above designed *in vitro* assay combined with structure analysis could act in high throughput workflow to find out the potential rules of adaptation on the level of this basis cellular function.

Since our results characterized the function of *HiFAS* in LA biosynthesis, the *in vivo* function of LA was still ambiguous. For LA and corresponding glycerides, the antibacterial effects have been characterized against a wide range of bacteria, showing relative high effect on Gram-positive bacteria (Borrelli et al., 2021; Casillas-Vargas et al., 2021; Hovorková et al., 2018; Suryati et al., 2023). With high LA content, raw fat extracts from BSF were tested *in vitro* to show positive antibacterial effects (Dabbou et al., 2020; Marusich et al., 2020). As addition in animal diets, BSF oil also exhibited positive effects in gut microbiota to enhance host immunity (Dabbou et al., 2020; Sypniewski et al., 2020). However, for BSF itself, cellular function of LA and its derivates have not been characterized. Due to knock down or knock out of *HiFAS* may cause growth defects as happened in other insects (Pei et al., 2019; Song et al., 2022), which may cover the consequential phenotypes of immunity impairment if LA and its derivates participate in BSF nonspecific immunity beyond its function in energy storage. Transgenic fruit fly loaded with HiFAS, act as *in vitro*, may detour these problems and reveal the biological function of LA in BSF.

## Author Contribution

YGJ and YPH conceived the study. YGJ designed research. YGJ performed all the experiments and analyzed the data. ZDP and PH assisted with fermentation. ZQK and BHC assisted with BSF rearing. YGJ and YPH wrote the draft. All the members commented on the manuscripts.

## Acknowledgements

This work was financially supported by the National Key Research and Development Program of China (XXXX). For material, the authors thank Dr. Yanlei Sun from CAS Center for Excellence in Molecular Plant Sciences for cDNA from fat body of *Drosophila melanogaster*. For sample preparation and detection, the authors thank Dr. Li Liu for freeze drying and Mr. Wenli Hu for GC-MS assistance in the Core Facility Centre of the Institute of Plant Physiology and Ecology.

