## Supplementary material for "Functional characterization of fatty acid synthase in *Hermetia illucens*": SM1.0.pdf

### Supplementary materials for Functional characterization of fatty acid synthase in *Hermetia illucens*

This profile includes:

Figure S1-5

Table S1-3

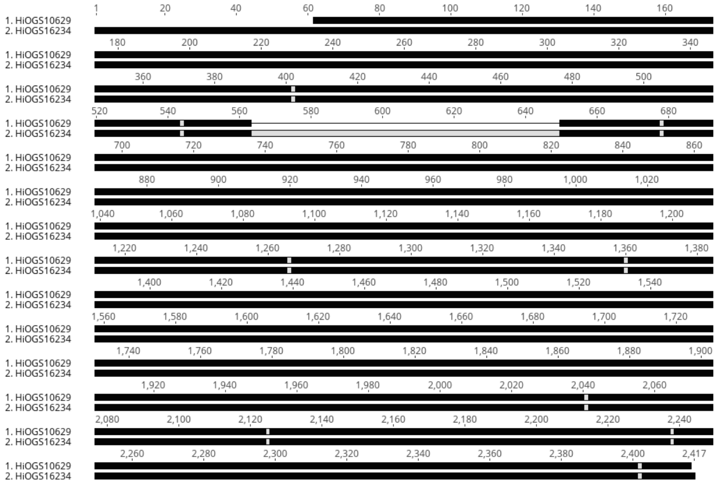


**Figure S1: Alignment of HiOGS16234 with HiOGS10629**

Alignment in protein sequences of HiOGS16234 and HiOGS10629 by identity. Despite several point mutations, compared with HiOGS16234, HiOGS10629 had two long indels: one from 1 to 61 in aa, the other from 564 to 649, which could not be proved its existence by molecular cloning.

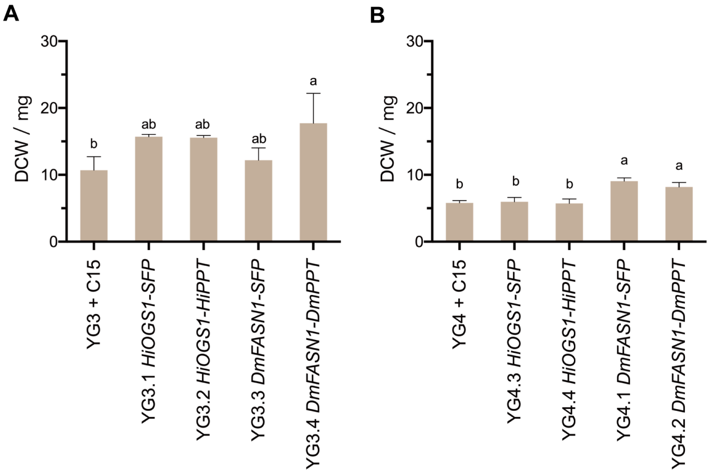


**Figure S2: Biomass accumulated in different strains after fermentation**

Dry cell weights were collected from different strains after vacuum freezing drying, including strain YG3 series (A) and strain YG4 series (B). The standard deviation was calculated from biological replicates. Different letters indicate statistically significant differences across groups using a one-way analysis of variance (ANOVA) with Tukey's multiple comparisons test, $P < 0.05$.

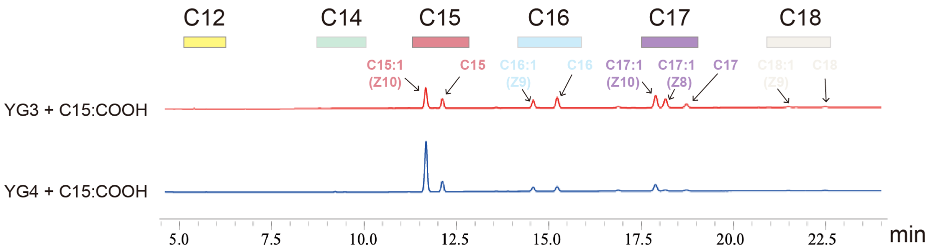


**Figure S3: Fatty acid profiles of *Scfas1*Δ strains before and after *ScELO1* deletion**

Gas-chromatography analysis of strain YG3 (*Scfas1*Δ) and strain YG4 (*Scfas1*Δ and *Scelo1*Δ). Obviously, further deletion of *ScELO1* in strain YG3 caused less elongation of pentadecanoic acid and more unsaturation of it.

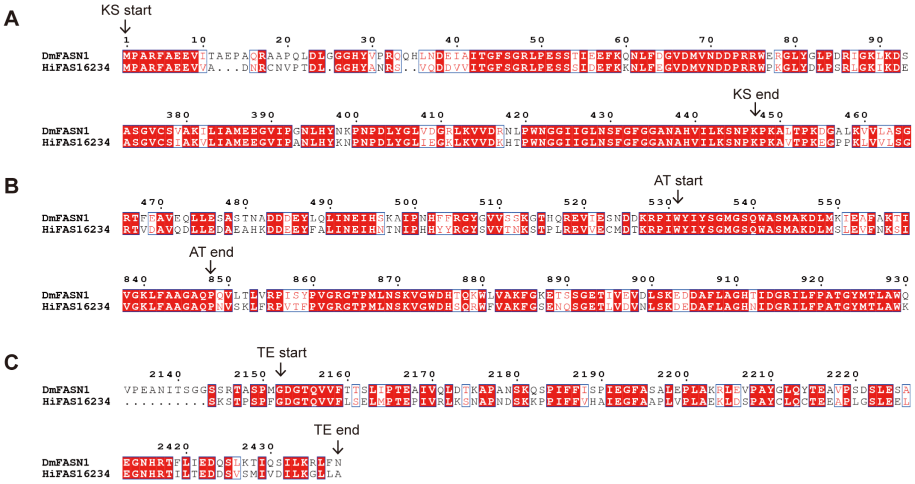


**Figure S4: Design of domain swap in HiOGS1**

Based on domain annotation by CD search in NCBI and protein sequence alignment, KS domain was identified from the $M^{1}$ to $K^{447}$ (A), AT domain was identified from the $W^{531}$ to $P^{848}$ (B), and TE domain was identified from the $G^{2152}$ to the last amino acid (C). The starts and ends of these domains were pointed by arrows.

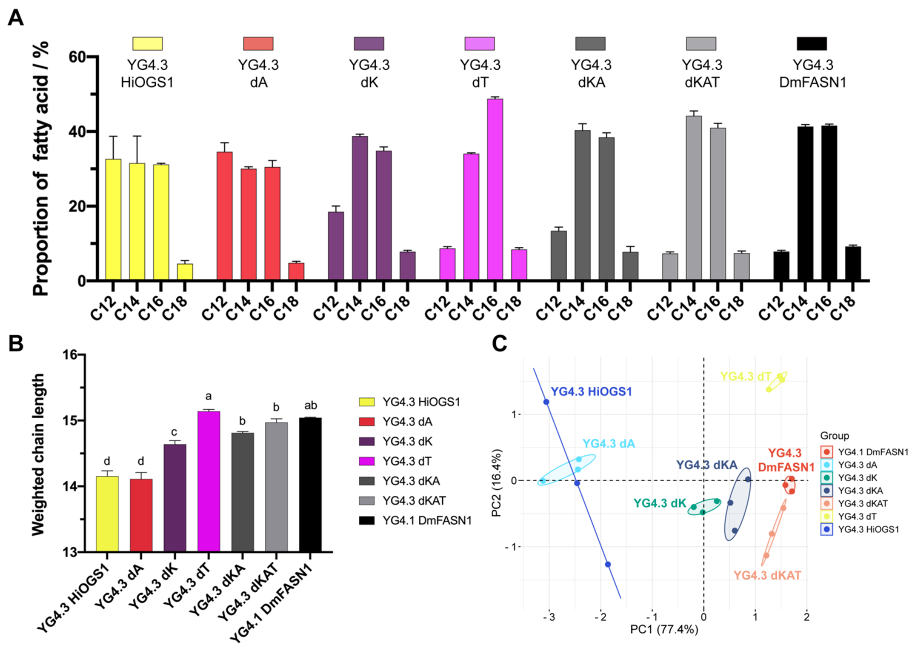


**Figure S5: Fatty acid profiles of chimera HiOGS1**

(A) Proportion of fatty acids in different chain length in strain YG4 series. (B) Weighted chain length of fatty acids in strain YG4 series. (C) Principal component analysis of fatty acid profiles in strain YG4 series. The standard deviation was calculated from biological replicates.

**Table S1: Primers for cloning and plasmids construction**

**Table S2: Strains used in this study**

| Strains | Genotypes | Resources |
|---|---|---|
| IMX581 | *MATa ura3-52 can1Δ::cas9-natNT2 TRP1 LEU2 HIS3* | Ref[1] |
| YG1 | IMX581 *pox1Δ faa4Δ faa1Δ gal80Δ* | Ref[2] |
| YG1.0 | YG1 *pESC-KlURA3* | This study |
| YG1.1 | YG1 *pESC-Gal1p-DmFASN1-Cyc1t-Gal10p-SFP-Adh1t* | This study |
| YG1.2 | YG1 *pESC-Gal1p-DmFASN1-Cyc1t-Gal10p-DmPPT-Adh1t* | This study |
| YG1.3 | YG1 *pESC-Gal1p-HiOGS16234-Cyc1t* | This study |
| YG1.4 | YG1 *pESC-Gal1p-HiOGS16234-Cyc1t-Gal10p-SFP-Adh1t* | This study |
| YG1.5 | YG1 *pESC-Gal1p-HiOGS07375-Cyc1t* | This study |
| YG1.6 | YG1 *pESC-Gal1p-HiOGS07375-Cyc1t-Gal10p-SFP-Adh1t* | This study |
| YG1.7 | YG1 *pESC-Gal1p-HiOGS16234-Cyc1t-Gal10p-HiPPT-Adh1t* | This study |

| | | |
|---|---|---|
| YG2 | IMX581 *gal80*Δ | This study |
| YG3 | YG2 *fas1*Δ | This study |
| YG3.1 | YG3 *pESC-Gal1p-HiOGS16234-Cyc1t-Gal10p-SFP-Adh1t* | This study |
| YG3.2 | YG3 *pESC-Gal1p-HiOGS16234-Cyc1t-Gal10p-HiPPT-Adh1t* | This study |
| YG3.3 | YG3 *pESC-Gal1p-DmFASN1-Cyc1t-Gal10p-SFP-Adh1t* | This study |
| YG3.4 | YG3 *pESC-Gal1p-DmFASN1-Cyc1t-Gal10p-DmPPT-Adh1t* | This study |
| YG4 | YG3 *elo1*Δ | This study |
| YG4.1 | YG4 *pESC-Gal1p-DmFASN1-Cyc1t-Gal10p-SFP-Adh1t* | This study |
| YG4.2 | YG4 *pESC-Gal1p-DmFASN1-Cyc1t-Gal10p-DmPPT-Adh1t* | This study |
| YG4.3 | YG4 *pESC-Gal1p-HiFASN16234-Cyc1t-Gal10p-SFP-Adh1t* | This study |
| YG4.4 | YG4 *pESC-Gal1p-HiFASN16234-Cyc1t-Gal10p-HiPPT-Adh1t* | This study |
| YG4.3 dA | YG4 *pESC-Gal1p-HiFASN16234 (DmAT)-Cyc1t-Gal10p-SFP-Adh1t* | This study |
| YG4.3 dK | YG4 *pESC-Gal1p-HiFASN16234 (DmKS)-Cyc1t-Gal10p-SFP-Adh1t* | This study |
| YG4.3 dT | YG4 *pESC-Gal1p-HiFASN16234 (DmTE)-Cyc1t-Gal10p-SFP-Adh1t* | This study |
| YG4.3 dKA | YG4 *pESC-Gal1p-HiFASN16234 (DmKS-AT)-Cyc1t-Gal10p-SFP-Adh1t* | This study |
| YG4.3 dKAT | YG4 *pESC-Gal1p-HiFASN16234 (DmKS-AT-TE)-Cyc1t-Gal10p-SFP-Adh1t* | This study |

#### Table S3: Sequences used in this study

>HiOGS16234

atgccggcccgttttgcagaagaagtagttgctgataatcggtgtaatgtccccacggacctagggggacattatgccaatcgatcggtacaa
gatgatgttgtcatcacgggtttctctggccgcttgccagaaagctcttcaattgatgagttcaagaagaatttattcgagggtgttgatatggttaa
tgacgatccacgaagatggcctaaaggtttgtatgacttaccatcacgtttgggaaagattaaagatgaagacttggaacaatttgatcaacaatt
cttcaacgtgcatcaaaagcaagcagagtgcatggacccacaattgcgtttgttgctagagactacacatgaggctattattgacgcaggagtt
aacccccaagatatccgaggatcccgaactggtgtctatattggagtgtcttcctctgaaactgaacaatactggtgcagtgatccggatttggtt
aacggttacggtttgaccggctgcgcgcgagctatgttcgccaaccgaatttcatttacattcgatttcaaaggaccaagtttctgcgttgacaca
gcctgctcgagttcgttgtatgctttggagcatgcattcgcagacattaaagccggaaaatgtgattctgcgatcgttgctggtgttggtctgatttt
gaagcctacaatgtccttgcaattcaaacgcctgaacatgttgagccctgatggtacatgccgagctttcgacgatagtggcgcaggatatgtg
cgatccgatggttgtgtgtcaatattccttcaacgtgctaaggatgccaaaaggatatactcgactatcctaaatgtacgaactaacaccgacgg
acaaaaggagcaaggtatcaccttccctgatggcaggatgcaaaaccgtttgattcgcgagacatacgaagaaattggattgaatccagcgg
aagttgtatacgttgaagctcacggcaccggaacaaaagttggtgatccacaagaggtaaactcaattacagactttttctgcaaggaccgtaa
gaccccactcttgattggatcagtcaaatccaacatgggtcactcagaacctgcatcaggtgtgtgctccattgcgaaggtccttattgccatgg
aggagggcgtcattccagccaacttgcattacaagaatccaaatccagatctctacggtctgattgaaggtaaactaaaagttgttgacaagca

cactccatggaacggtggcatcatcggcttgaattcgttcggatttggaggtgccaacgcccacgttatcttgaaatcgaatccaaaacccaag
gcggtgacccctaaggaaggcccaccaaagctagttgttttatccggtcgaacagtagatgccgttcaagatctgcttgaagatgctgaggca
cataaggacgacgaagaatatttcgcactcatcaacgaaatccacaacacaaacatcccacaccattactaccgaggatactctgtagttacaa
acaagtcaacccccttgcgtgaagtcgtggaatgtatggacacaaagcggccaatttggtatatctactcaggaatgggaagtcaatgggcta
gcatggcgaaggacttgatgagtttggaagttttcaacaaaagcattcaccgatgcgctgaagtgctccgtcctgagggtgttgatcttattgata
tcttaacccgctcgaatgaggctcgctttgacaacatcttgaactcatttgtgtcaatcgccgccatgcaagttgctctaactgatcttctgaccag
cttaaatatttctccggacggcatccttggacattccgtcggtgaattgggttgtgcctatgccgatggttgctttaccccagagcaaacagtact
ggccgcctactggagaggaaagagcatccaggacactgatctcgccaagggtcagatggcagccgttggtttatcctgggaagagtgcaaa
caacaactgcctgagggcgtgattgctgcttgccacaatagttcagacagtgttacaatctccggaccagaaggcaaggttactgaagtcgttg
agaagctttcagccgctggtgtgttcgccaaggctgtcaaatcctcgggctatgcattccacagcaaatacattgccgatgctggtccaaaact
acgcaagagcttggaaaagatcattccgaacccgaaaaatcgtacctcccgttggattagttcgagcataccagaaagtgcatggaatactcc
aattgctaaacagagctctgctgcttaccatgtgaacaacttgctttcgccagtcttgttccacgagggtcttcaacatgtgccaaagaacgctatt
tgcatcgaaatcgcaccacatggattgttgcaggctattctgaagcgagctcttggtggaggtgcaacaagcttaagcttgattaagcgctacc
atgacaacaatctggaattcctcttaacaaacgttggaaagttattcgcagccggcgcccaaccaaacgtctcaaaactcttccgtcctgttacat
tccctgtaggccgcggcactcctatgttgaactccaaagtaggatgggaccattcacagagatggtttgtcgcgaaattcggctcagaaaatca
atcgggcgaaacccttgtcgatgtcaacctatccaaggatgaggatgctttccttgccggtcacaacatcgatggacgtatcttgttcccggcta
caggctatatgaccctggcatggaagactttcgccaaggttcactccagtgacatgtcacaaacccccgtagttttagaaaacgttgtcttccat
cgtgcaacaattttgaataaggataatgctgtgaaatttggtatcaacttcttcgatggtactggcaactttgaaatttgcgaaggaggttcactgg
cggtgtctggtaaaatcacaattccggatcatgtagacaaggaggagcttccattgaatccattggaaaaagacaaatcaggactaccgttga
gcaagaacgatgtatacaaagaattgagacttcgaggctatgattacagtggcatgttccgaggagttaaggaaaccgacaaccaagcagtta
caggaagcttggattgggaaaacaactgggtcagtttcatggacacaatgctgcaattcagcattatgggcaaggacttacgagagctctatct
tccaacacgaattgagaagatcatcatcaacccagcaaaacacctcgaacttcttgcacaagtttcggacgagaagaaatcactccctgtatac
atgtacaaggacattaatgtgatcaagagtggtggtgttgaaatgcgtggcttgaaggcttcccttgcaccacgccgttccggcactcaactcc
caccctctttggagcgctacgtcttcgtaccacaaagcaatggctccgatctatctgaaaaccctgagaaggcacgcttgaacgccctcaccgt
ggcattgcacttagtcactgataattctggcggcgcactcaaagttaaagttgctgaaattgcaaacgaacgtaatgctgacagcttattggccc
aaaacattcttggcataattgaaggcgagcccaccttagcctccgatgttgttgtggcttccgctaaaaataccgatcaactgaaatctcactttg
gagattctggagtccgcgtagttaacaagaacatttctgagggaccagtcgaacaaaattgccatcttgtgctcgcatacgatgttcttggccgt
ccagatagtgcaaccgttctggaaaacgtgctgaacacaatcagagaggaaggattcattcttttggatgaagtccaatccaactacgaaaag
aacgcttccgtcctacaaaagaagaatttgattgtggtttccacccaaaaggtcggccaacgagtatttgtattggtccgctcgcttatcgatgtc
cgcaacaggaagaaggtcgtcattcagttgacggaaaagaacttcaagtggttggatcaagttaagaaagaattgcaaacagccgaagatga
gaacaaattcatttatttagtctgccaaggcgaggaattgtttggcgctgttggtctcctgaattgcgtcaagaatgaagtaggcggtcgattcgc
aaggatgtacttcatccagaatccatcagctgagaaattctcagtggattctaaattctaccaggagcagctgaataaggatcttatttgcaatgta
ttgaagtcggccggctggggaaccttccgtcacttgaaattagagacgtctggagacacccctactctacaagtcgaacacgcttacatcaatg

ctatcaccaaaggtgatttgtccagtctcaagtggatcgaaggacctctctcagtccagaaaccggcatttggagataaccaggaagatctctg
cactgtctactatgctccaatcaacttccgtgacgtcatgctgagctctggaaaattagctgccgatgcattacctggagatttagcacaacagg
actgtgtcctcggtttggagtttgccggccgtgactcaagcggcaaacgcatcatggccatggttacagccaaatcattggccacaacgtgcg
ttgcttcccgcaacatgatgtgggaaattccagacaattggacaatggaacaagcctcgacggtgccttgtgtctactcaacagtgtactacgct
cttgtagtgcgcggtaaaatgaagaagggcgagaagattctcattcacgctggtacaggtggtgtcggacaagccgcaatctccgttgcattg
catcatggattgactgtattcactactgttggaagtcaagagaagagagacttcttgaaaaagactttcccacaattgactgaccgaaacattggt
aactctcgtgatacatcctttgaacagttgatcatgcgcgaaacacaaggtcgtggagttgatttggtactcaactcgttggctgatgaaaaactg
caagcctccgtcagatgtttgggattgaatggacgattcttggaaattggaaaattcgatttgagcaacaactcgccgttgggtatgtctgtcttct
tgaagaacacatcgttccatggaattttgttggacagcatcatggaaggtgacgaagaaacacagaaagaagttgtggctttagttgcggagg
gtatcaagaacggtgccgtgcgtcctctaccaacaactgtcttcaatgaacaacaagtagagcaagcattccgtttcatggcttctggcaagcat
attggcaaagtcgttattaaagtacgtgatgaggaatcagctaaggttctgaaaccacaaactaaattggtgaccgctatcccaagaacatacat
gcatccagagaagacatacattgttgtgggaggattgggtggcttcggattggaattgtcccattggatggtgtcacgtggtgctaagaagctg
gttctcactagcagaagtggtgttaagaccggctatcaatctctgatgatccgtaggtggaaggcccgcggtgtgcaagtcgccattgacacc
aatgatgtaacgacatctgaaggatgcaaaaaactacttgaagcttcgaataaacttggattggttggcggaattttcaatttggcggcggtact
gagggacggactacttgagaaccaaaccgaacaagactttaagactgtatgcgttcctaaggtggatggtaccaaatacttagatcaattgtca
cgagccttgtgtcctgaattggattacttcgtctgcttctcaagtgtgtcttgcggtcgtggtaacatcggtcaaacaaattacggcttggctaact
cagccatggaacgaatttgcgaaaatcgtcaagcatctggcttcccaggaactgcaattcaatggggtgctatcggtgatactggtcttgttatt
gagaatttgggcgacaacgacactgttattgggggaaccctaccacaacgaatgacatcatgtctccaaacgttcgacatgttcctacaacaac
ctcacgctgtattggcatcaatggtcgtggctgaaaagaggaaggctgacgcaactggaggagtcagtttagtaaatagcattgcaaatatcct
aggcttacgagacatgaggaacgttcaagacgttgcaaccttagctgatctcggaatggactctctcatgggtgctgaaatcaaacaaactctt
gaacgaaactttgatgttgtgatgacaccacaagaaattcgtcaattaacattcggcgcacttaaggagctcgagggttctgcaggcggaacc
gtttcaccagctccagtcagtccatcaaagagcaccccatcaccattcggcgatggaacccaagtagttttcctatcagagctgatgcccaccg
aaccaattgttcgtttgaaatcaaacgcaccgaatgactcaaagaaacctccaatcttcttcgtgcatgccattgaaggatttgccgcacctttagt
accattagccgaaaaaattagacagtcccgcctattgtttgcaatgcacagaagaggcaccacttggatcacttgaggaattaggcgaatattat
gttaaacaaattaagaaaatccaaccgaaaggaccttactcgatagctggctattcgtttggtgcaactattgcttatgtaatggcagcttacttag
aacaacagaatgaggatgtacaacttgtgatgttggacggtgcacccaagtatgtatcgtggtacacacaaacacataaggcaaagaaaaat
gcattggaatcaacagaggaacagaatgaggcttatgcgttggcatactttggcatggttctaggaaatatggattatggtggtgttgctcgttta
cttcttaaaatcccaacatgggaggccaaggtcgaaaaatgtgcagaattgcttcatgcagaaattaatcaaccagttgacttgattatcaaaact
gcaactgcattctaccacaaactcttggctgctgacaaattcagagcatcacataagatcaagaagtctgtaacgcttatcaaacccactgaaaa
ttatgccaaactagaagctgattacggtctttcggaggtatgtcaaaaggatgttaaaatcctaacagtcgaaggaaaccatcgaaccatcctaa
cagaggatgattccgttagcatgatagtagacatcctcaaaggactcttagcataa

>HiOGS10629

atggttaatgacgatccacgaagatggcctaaaggtttgtatgacttaccatcacgtttgggaaagattaaagatgaagacttggaacaatttgat

caacaattcttcaacgtgcatcaaaagcaagcagagtgcatggacccacaattacgtttgttgctagagactacacatgaggctattattgacgc

aggagttaacccccaagacatccgaggatcccgaactggtgtctatattggagtctcatcctctgaaactgaacaatactggtgcagtgacccg

gatttggttaacggttacggtttgaccggctgcgcgcgagctatgttcgccaaccgaatttcatttacattcgatttcaaaggaccaagtttctgcg

ttgatacagcctgctcgagttcgttgtatgctttggagcatgcattcgcagacattaaagccggaaaatgtgattctgcaattgttgctggtgttgg

tctgattttgaagcctacaatgtccttgcaattcaaacgcctgaacatgttgagccctgatggtacatgccgagctttcgacgatagtggtgcagg

atacgtgcgatccgatggttgtgtgtcaatattccttcaacgtgctaaggatgccaaaaggatatactcgactatcctaaatgtacgaactaacac

cgacggacaaaaggagcaaggtatcaccttccctgatggcaggatgcaaaaccgtttgattcgcgagacatacgaagaaattggattgaatc

cagcggaagttgtatacgttgaagctcacggtactggaacaaaagttggtgatccacaggaggtgaactcaattacagacttcttctgcaagga

ccgtaagaccccactgttgattggatcagtcaaatccaacatgggtcactcagaacctgcatcaggtgtgtgctccattgcgaaggtccttattg

ccatggaggagggcgtcattccagccaacttgcattacaagaatccaaatccagatctctatggtctgtttgaaggtaaactaaaagttgttgac

aagcacactccatggaacggtggcatcatcggcttgaattcgttcggatttggaggtgccaacgcccacgttatcttgaaatcgaatccaaaac

ccaaggcagtgacccctaaagaaggcccaccaaaactagttgttttatccggtcgaacagtagatgccgttcaagatctgcttgaagatgctga

ggcacataaggacgacgaagaatatttcgcgctcatcaatgaaatccacaacacaaacatcccacaccattactaccgaggatactccgtagt

tacaaacaagtcaactcccttgcgtgaagtcgtggaatgtatggacacaaaacgaccaatttggtacatctactcaggaatgggaagccaatg

ggctagcatggcgaaggacttgatgggtttggaagttttcaacaaaagcattcaccgatgcgctgaagtgctccgtcctgagggtggtcagat

ggcagccgttggattatcctgggaagagtgtaaacaacagctgcctgagggcgtgattgctgcttgccacaatagtgcagacagtgttacaat

ctctggaccagaaggaaaggttactgaagtcgttgagaagctttcagccgctggtgtgttcgctaaggctgttaaatcttcgggctatgcattcc

acagcaaatacattgccgatgctggtccaaaacttcgcaagagcttggaaaagatcattccaaacccaaaaaatcgtacctcccgttggattag

ttcgagcataccagaaagtgcatggaacactccaattgctaaacagagctctgccgcttatcatgtgaacaacttgctttcgccagtcttgttcca

cgagggtcttcaacatgtgccaaagaacgctatttgcatcgaaatcgcaccacatggattgttgcaggctattctgaagcgagctcttggtgga

ggtgcaacaagcttaagcttgattaagcgctaccatgacaacaatctggaattcctcttaacaaacgttggaaagttattcgcagccggcgccc

aaccaaacgtctcaaaactcttccgtcctgttacattccctgtaggtcgcggcactcctatgttgaactccaaagtaggatgggaccattcacag

agatggtttgtcgcgaaattcggctcagaaaatcaatcgggcgaaacccttgtcgatgtcaacctatccaaggatgaggatgctttccttgccg

gtcacaacatcgatggacgtatcttgttcccggctacaggctatatgactctggcatggaagactttcgccaaggttcactccagtgacatgtca

caaacccccgtagtgttagaaaacgttgtcttccatcgtgcaacaattttgaataaggataatgctgtgaagtttggtatcaacttcttcgatggtac

tggcaactttgaaatttgcgaaggaggttcactggcggtgtctggtaaaatcacaattccggatcatgtagacaaggaggagcttccattgaatc

cattggaaaaagacaaatcaggactaccgttgagcaagaacgatgtatacaaagaattgagacttcgaggctatgattacagtggcatgttcc

gaggagttaaggaaaccgacaaccaagcagttacaggaagcttggattgggaaaacaactgggtcagtttcatggacacaatgctccaattc

agcattatgggcaaggacttacgagagctctatcttccaacacgaattgagaagatcatcatcaacccagcaaaacacctcgaacttcttgcac

aagtttcggacgagaagaaatcactccctgtatacatgtacaaggacattaatgtgatcaagagtggtggtgttgaaatgcgtggcttgaaggc

ttcccttgcaccacgccgttccggcactcagctcccaccctctttggagcgctacgtcttcgtaccacaaagcaatggctccgatctatctgaaa

accctgagaaggcgcgcttgaacgccctcaccgtggcattacacttagtcactgataattctggcggcgcactcaaagttaaagttgctgaaat tgcaaacgaacgtaatgctgacagcttattggcccaaaacattcttggcataattgaaggcgagcccaccttagcctccgatgttgttgtggctt

ctgctaaaaataccgatcaactgaaatctcactttggagattctggagtccgcgtagtcaacaagaacatttctgagggaccagtcgaacaaaa

ttgtcatcttgtgctcgcatacgatgttcttggccgtccagatagtggaaccgttcttgaaaacgtgctgaacacaatcagggaggaaggattca

ttcttttggatgaagtccaatccaactacgaaaagaacgcttccgtcctacaaaagaagaacttgattgtggtctccacccaaaaggtcggcca

acgagtatttgtattagtccgctcgcttattgatgtccgcaacaggaagaaggtcgtcattcagttgacggaaaagaacttcaagtggttggatc

aagttaagaaagaattgcaaacagccgaagatgagaacaaattcgtttatttagtctgccaaggcgaggaattgtttggcgctgttggtctcctg

aattgcgtgaagaatgaagtaggcggtcgattcgcaaggatgtacttcatccagaatccatcagctgagaaattctcagtggattctaaattcta

ccaggagcagctgaataaggatcttatttgcaatgtattgaagtcggccggttggggaacattccgtcacttgaaattagagacgtctggagac

actcctactctacaagtcgaacacgcttacatcaatgctatcaccaaaggtgatttgtccagtctcaagtggatcgaaggacctctctcagtcca

gaaaccggcatttggagataaccaggaagatctctgcactgtctactatgctccaatcaacttccgtgacgtcatgctgagctctggaaaattag

ctgccgatgcattacctggagatttagcacaacaggactgtgtcctcggtttggagtttgccggccgtgactcaagcggcaaacgcatcatgg

ccatggttacagccaagtcattggccacaacttgcgttgcttctcgcaacatgatgtgggaaattccagacaattggacaatggagcaagcctc

tacggtgccttgtgtctactcaacggtgtactacgctcttgtagtgcgcggtaaaatgaagaagggcgagaagattctcattcacgctggtaca

ggtggtgtcggacaagccgcaatctccgttgcattgcatcatggattgactgtattcactactgtcggaagtcaagaaaagagagacttcttgaa

aaagactttcccacaattgactgaccgaaacattggtaactctcgtgatacatcctttgaacagttgatcatgcgcgaaacacaaggtcgtggag

ttgatttggtactcaactcattggctgatgagaaactgcaagcctccgtcagatgtttgggattgaatggacgattcttggaaattggaaaattcga

tttaagcaacaactcgccgttgggtatgtctgtcttcttgaagaacacatcgttccatggaattttgttggacagcatcatggaaggtgacgaaga

gacacagaaagaagttgtggctttagttgcggagggtatcaagaacggtgccgtgcgtcctctaccaacaactgtcttcaatgaacaacaggt

agagcaagcattccgtttcatggcttctggcaagcacattggcaaagtcgtcattaaagtacgtgatgaggaatcagctaaggttctgaagcca

caaactaaactggtgaccgctatcccaagaacatacatgcatccagagaagacatacattgttgtaggaggattgggtggtttcggattggaat

tgtcccattggatggtgtcacgtggtgctaagaagctggttctcactagcagaagtggtgttaagaccggctatcaatctctgatgatccgtagg

tggaaggcccgcggtgtgcaagtcgccattgacaccaatgatgtaacgacatctgaaggatgcaagaaactacttgaagcttcgaataaactt

ggattggttggcggaattttcaatttggcggcggtactgagggacggactacttgagaaccaaaccgaacaagactttaagactgtatgcgttc

ctaaggtggatggtaccaaatacttagatcaattgtcacgagccttgtgtcctgaattggattacttcgtctgcttctcaagtgtgtcttgcggtcgt

ggtaacatcggtcaaacaaattacggcttggctaactcagccatggaacgaatttgcgaaaatcgtcaagcatctggcttcccaggaactgca

attcaatggggtgctatcggtgatactggtctcgttattgagaatttgggcgacaacgacactgttattggtggaaccctaccacaacgaatgac

atcatgcctccaaacgttcgacatgttcctacagcaacctcacgctgtattggcatcaatggtcgtggctgaaaagaggaaggctgacgcaact

ggaggaatcagtttagtaaatagcatcgcaaatatcctaggcttacgagacatgaggaacgttcaagacgttgcaaccttagccgatctcgga

atggactctctcatgggtgctgaaattaaacaaactcttgaacgaaactttgatgttgtgatgacaccacaagaaattcgtcaattaacattcggc

gcacttaaggagctcgagggttctgcaggcggaaccgtttcaccagctccagtcagtccatcaaagagcagcccatcaccattcggcgatgg

aacacaggtagttttcctatcagaactgatgcccaccgaaccaattgttcgtttgaaatcaaacgcaccgaatgactcaaagaaacctccaatct

tcttcgtacatgccattgaaggatttgccgcacctttagtaccattagccgaaaaattagacagtcccgcctattgtttgcaatgcacagaagagg

caccacttggatcacttgaggaattaggcgaatattatgttaaacaaattaagaaaatccaaccgaaaggaccttactcgatagctggctattcgt

ttggtgcaactattgcttatgtaatggcaacttacttagaacaacagaatgaggatgtacaacttgtgatgttggacggtgcacccaagtatgtat
cgtggtacacacaaacacataaggccaagaaaaatgcattggaatcaacagaggaacagaatgaggcttatgcgttggcatactttggcatg
gttctaggaaatatggattatggtggtgttgctcgtttacttcttaaaatcccaacatgggaagccaaggtagagaaatgtgcagaattgctgcat
gcagaaattaatcaaccagttgacttgattatcaaaactgcaactgcattctaccacaaactcttggctgctgacaaattcagagcatcacataag
atcaagaagtctgtaacgcttatcaaacccactgaaaattatgccaaactagaagctgattacggtctttcggaggtatgtcaaaaggatgttaa
aatcctaacagtcgaaggaaaccatcgaaccatcctaacagaggatgaatccgttagcatgatagtagacatcctcaaaggactcttagcataa

>HiOGS07375
atggcaccaaatgagaattctacttcagcggctgttcaaaatggataccataagcgaacctcaaggatttttccagaaaccgatgaagacgata
ttgtcgtatctggaatttcaggaagatttcctaactctagaagcattagtgaatacgaatacaatctctataacaagattgacatggtagatggcga
agagacccgctggaaaagctttaacccggaaatccctcaaagaggtggaaaaatctacgatttggagaagtttgacgctactttcttcggggta
catttcaagcaagctcacactttggatcctcaatgtcgaatcttgatggaagtagcatatgaagctatattggacgcaggagtcaatcctaaaagt
ttaaggggcacgagaactggagtatatgtgggcggatgcatatcggaatcagaaaaaacatggttctacgaaaaaatttccactggaggattt
ggtattaccggttgcagtcgtgctatgttagcaaacagaatttcttacgtattggggctcaatgggccttcatatgtagtagataccgcttgcagta
gttcaatgtatgccttagacaatgccttctctgcttttcgtaacggtgaaattgatgctgcaatcattggtggagttaacttaacccttcatccctatgt
tactttgcaattcgcccgattgggcgttttagcagctgatggatattgtcgaccatttgatcaagatgccagtggttatacccgttctgagtcggtg
aattgcttgtttttgcaaagaaaacgtgatgcaaagcgtatctatggctctctggtctactcaaaaaccaattgcgatgggttcaaaacagaaggc
ataacatatccttcaggaatagtacaagagaaactacttcgcgaattttataacgaaatagggatcagcccagataaactgggatatttggaagc
gcatagtacaggaactttagttggcgatccagaagaatgtcgcgctattgatacggtcttatgcaaatctagaagcaaacccctacttgttggat
ctgttaaatctaatattggacactcggaagcagcttcaggtatatgttccatggttaaggccatattggccttagaaaaaggcgagattgccccg
aacatcaatttcaaaagtgttcgtaaagaaattccatcattgctagaaggaagattgaaggttgtaagtgagatcgaaaaactcaccactccctat
attggagttaattcatttggatttgggggtgccaatgctcacgctcttctcaaaggctgtgaaaaggaaaaacaaaatcagggagcacctgaag
atgacttgccgcgtcttgtgtattggagcagtcgcactgaagaaggcgtgaatgtaatgctagattctatagagtccaatccattggatactgagt
atcttgctctaacgtataatatacaaaaagaagggattccagggttcgtattcaggggatatggtttatattctcacgtgaaaggaggaaatgcta
cttgtatatcgaaagaagtgcaacattacactggagttcaaagaccaattgtttgggttttcagtggtatgggatcccaatggacagagatggga
tcttcgctcatgaatataccaatgttccgtgaatccatcaatcgatgccataatgcactggcacctaaaggattaaatttaattaatatcattacatct
tcagatccgaaaatgtttgataatatcctaaattcatttgtgggcatcgcggctattcaaatcgcgatcgttgatattctaagagcactagattttga
acctgattatcttattggtcattccgttggtgagctaggttgtgcatacgctgatggaggattcactgctgaacaaatgattctatctgcatactctc
gtggaaaagtcagcctagaagttgataaaattaagggttcaatggctgctgtgggtctcggttacaaaaacatcatcaaactattaccccccttca
attgaggtcgcatgtcataacagcgctgactcgtcaacaatatctggcccagcagaagatgtggaaaagttcgtagcagaactgaaaagtaaa
aatattttcgctaaggaagtaccatgctcaaatattgcatatcactctcgctatattgcccacatgggaccgctgcttcttaaatatctccaagaaat
tattccgaatccaaaaaaacgttcacaaaaatggataagttcgagcgttccaaaacaccaatgggatcaacctcgcagtgaatattcttctgccg
aatatcataccaataatttgttgaatagtgtactctttgaagaagcagcaatgcttctacctccaaatgcactgacggttgagattgctccacatgg

attgttgcaggctattttaaagaaatcaatgccaaatgctgttaatatcggtctcacccaacgtgggaacaaaaataatgctaacttcttcttgtctg

ctttaggaaaaatccatttaaatggattcactttacccatcgaacgattgtatccgaaagtggaatttcccgtctcacgcggtacgaaaaccatttc

ttctctaattcgttgggatcacggtgaaaattggtttgtatccaaatttgaaaacatcagatctgaaaagacaggcgaaagggtattccaaattgg

cttgaatgctgaagacaccgaattcctggctggacatatcatcgatggaaaggtcctacttccagcaacattctacctttctatggtgtgggagg

cgttctcattgatgcacgctggacctatgtacatgaatatgcccgttgaattcaatgacgtacgtttcgttcgagctacacatttaaacaagggttc

aaatgtaaccttaacagtgtcaatacatcacggaacaggtctattcgatatcactgaagccggagcgatagtagtgaccggaagaataaatga

aattatgaatccgcaacctccggagccattgccaccatgtccatctaacgactgcactatgcttgaaacgcaagatatctataaagaactcagat

tgcgaggatatcattatgctggagcattcaagagtgttatgcaagctcgtagtgatggggtcactgggaaagtcaaatggtcattcaactgggtt

gctttcatggattgtctcctgcaaatacatatccttggatctgattcgcgttcattaatacttccaacccgtatccggcagcttcgcattcttgggctg

caccacattaatttagcagcgcaaatggacccgcagaacaaatacttcgatgtccagatggatcatactacacagcggttagttgctggtggga

ttgaaattttggggcttcaagccagtccagtttctcggagaaaaccaccaggtattccagttttggaaacctacaaattcatcccgcatttgccag

ctcccacaactgaaataactgacgctgcaaggatttgcgtgcaaactgctttggaaaataatcccgtaccaaaaatgaaaattgtagaggtgga

tcaagataaggttccaccaactattccgcacttccaagaggcattagaagacttgccggtggttactggcgacttcatgcttttaacttctcaaga

attagaactgcctgatattcacgttgaagatggaaaattgtcgactcaaacaaactgcacatttgtcatatccacaaacctcttgggtcgaccaga

atgggtccagaaggcccaaaagtcaattggcgatggtggatacttagtatcaatcgaaactcttcctctcaactatgcacaagtcaaagcgcct

ataggattcaatattattgcatttttaccaaccgaaaagggatcggtggtccttatgagacgacatccaaaaaaagtaacaaatatcactgttatcc

atgttgttcaaagcgatatgaactacacctggatcgaaaaaattaaaaatgcacttgctattaacccagtgattttagtagcccagaatgagaacc

tgaatggtctaattggcttgacgaattgcttacggaaggaaccaagtggacatcaaattacctgcgtgtttattgacgatcctcaagcaccagag

tttgacctagaacatcccttctatgctgataaattgaagttcggactgggaatcaatgtttttaagaatggtcaatggggaacatatcgacatttact

actgaagccattcctagaacccaaaaagaataaacgacaattatacgttgaccagcttcaacgtggtgatctatcatcacttacttggttcgaag

gacgaaagtcagtcgaagattgccaaatcaaagtagtgtatgcctccctcaacttccgagatatcatgctcgcaactggtcgaatcacattaga

atcacaccagagctcacgtttggaaaaccaatgtctgcttggattcgagttcgcgggtgtaggaaaagatggaaaacgatatatgggattgaa

gagcaatagaggtctggctacacatatcgatgaaactcagatcttgtggaacgtaccagatgattggacattggaacaagcggctacaattcct

tgcgtttatgttacggtttacatggctttcttccttgtgagcgatatccgtaagggtaagtcaattcttatccacgctggaacaggaggagttggac

aagcagctatcactgtggctttggctcatggacttgaagttttcacaaccgttagtacaaaagagaaaaagcagttcctgcttcaaaagtttccaa

aattaaaggaatcgaatatcggtaattctcgtgatacttcattcgaatatatggttctgaggcaaacaaaaggaaaaggagtggatttcgttctca

actcattggcggaagataaactgctagcttctgtacgctgtttaggaaataaggggcactttttcgaaattggaaagtttgatattatgaacgaca

gcaagctcggcttagcagtctttgaaaaggaaatatccttccatgcgatatttgctgatcgcttattcttcgcaggaatgaatgtcattaaagatgtt

atggagttgatctataagggcatccgtgaaggtgttataaaaccattgcctgtaacagtgttcaaagcagatcaagttgaacaagctttccgtcat

atggcagctggaaagcatatcggtaaagttctaattcaagttcatgctgagaacgataaggagcacaatacaatagtaccaatatcacctcttcc

gcaaacttatttcgatcgcaatatggtttatataattcctggaggccttggaggatttggtttggaactggctgattggatgattttgagaggagctc

gaaagttggtactcagttccagtaagggtttttcgaaaccatatcagccgtacagggtgaaggtatggaaatcatatggagccagagttgaaatt

agaactgatgatataaccacctatgaggggtgcctgtcattaataaaagaatcaatgaaaatgggggccatcggcggcattttcaatctagccg

ttgctttacgtgatgcgatcttccttaatcaaacaccagaaaagttccaagaatgtttagcaccaaaagctgtggccacaaaatatttggatgaaa
tatcacgaaaactgtgtcctcgtttgaaatatttcgtagtattttccagcgtgtcttgtggtcgaggtaacgctggccaaacgaattatggtatggc
gaattctataatggagaggattatcgaacagcgcgccctcaaaggcttgccagcgaaagctatacaatggggagctgttggagaagttggtct
cgtagccgacatggctgaagataaaattgatatggaaataggcggcactttgcaacaaagaatctcatcctgcctacaagagttggacaatcta
ttatccacgaatgatgcgattgtatctagtatggttgttgctgagaagcggcgcataaagagtggaaacataatggaaactgttctaagtattatgt
ctatacgtgacttgaagtcagtttcaatgggcacctcgctttccgaaactggtatggattccttaatggctgtcgaaattaaacaaactttggaaag
agagtttgaattgttcctcacacctcaagatttgcgaacaatgacattccagagattacaagaaatttcagatgctaaattgaaggatgaaggag
agcaggtgaagttacgacttgcacgtgatgataaaccggttggaatggctctgctacttcgaaatctcggggatgaattgaatagtcaccatac
catactgcctctgattacgaagaatagtaacctggtggatttgaattcgtcgcttatcattcctggcctagaaggtgtagctggacaagcgtggta
caatgtcggtctaggactttattctccttcgtatatcttgcaactcatgggcaccgtacatttatcatccatacaggaaatcgccgcttctgtgtttga
gaaagcctcacaagtatttaagagaaatcagcccttctacctggtaggctactccttcggtacgtttattacactggaactagcatacatgctgga
acaattaggttttcaaggatatgtaacaattatcgatggttcaccattatttttgaaaaaactttccctgcggacctttaaagaatcttcctcggacga
tgagattcaatctatgcttcttactacaacgctcatgcaaatatttcctggagagtttgacgaagatactacagttaaatttttggcgctcccggatt
ggaacagtcgcattgagaagtacttagaatatgcagaaaaacaaactgtttatagcagagactatgctgagtgtataatgcgagcaatgtttgat
cgtatcaaaatggtcaatgatctttcgttgaatgacgttaaacgcatccaatcatcaattacattggtccgacccacggatgtgtcagttcccgata
tcgatgaggattatgagctgtgtaaatacactactggaaaagttacagttaagtttattgaaggtgatcacacttccatgctggacaatcctaagtt
accgcaaataatcaacagcttcgatccgaatcttgaatctgataaatgttttgaagatctattataa

>HiPPT

Atgctgccgaaatacagttgtgtgcgctgggcgttagatttatcacagtggcgcccgacacttgccgagctcagcttagcgactgcatgtttac
aagccgaggagaaggctcgtctcatgaaattctacttccgggatgatttcgatgcatccttaatagggcgcctgctcatgcggaaatttgtgcg
ggacacaaccggaatcgcttacgatcaaataagattcgagcgtgatggacgcggcaaacccttcctggtcaatagcgtcttgggaccgtgtc
ggatcgacttcaacgtttcccatcagggcaactatagcgtgctggccggaatcgttgagagatcaattgcagacgagccacctgtgaaaatcg
gaatcgatgttatgaaaattgagtattcaggtggaaagccattaaccgaattcttccgcctgatgacgaggaacttttccgacgccgagtggag
atttatccggagccgacagtctgaggccgagcaattaaaagcgtttatgcgattctggtgtctgaaggagagttacgtcaaaaatgtcggtgttg
ggattaccatgaatttacagaaaataagtttcaaggtgaagactgacgacgtagcgaaagggaaggtggttcaggacacggtcttggaggtc
gatgggaatttactgacgaattggtgttttgaggagcatctgctggacgagcagcacagcgtagccatagcatttgaaaatttaggaaaaatgg
cagatgaaggcacatttaaagttttaggatatgaagagttgatggaaggcgcattgcccctaatggaaaaagataattcttattgtcaaagggttt
tggaaaagcaatataaacgtttccaataa

>DmFASN1

atgcccgcccgattcgccgaggaagtcatcaccgctgagcccgcacaacgtgccgccccccaattggacctgggtggtggtcactatgtgc
cccgtcagcagcacctcaacgatgagatcgccatcaccggtttctccggccgtctgccggagagctccaccatcgaggagttcaagcagaat

cttttcgacggcgtcgacatggtcaacgatgatccccgccgttgggagcgtggtctgtatggacttcccgacaggattggcaagctcaaggat
agcgatctggagaactttgaccaacagttcttcggtgtgcaccaaaagcaggctgagtgcatggacccactgctgcgcatgctgctcgagctc
acccatgaagctattattgacgctggcctaaacccaagtgatctgcgcggttcccgcactggtgtgtacatcggtgtgtccaactcggagactg
agcagcactggtgcagcgatgctgaccgggttaacggctacggcttgacaggatgtgcccgtgccatgtttgccaaccgcatctccttcacct
ttgatttcaagggacccagctacagcattgacaccgcttgctccagttctctgtacgccttggaacaggctttctccgatatgcgcgaaggaaag
gtcgacaacgccctggtcgctggagctggactgatcctcaagcccaccatgtcgctgcagttcaagcgactgaatatgttgagcccggacgg
cagctgcaaggccttcgatgagtctggcaatggatacgtccgttccgatggatgtgtggtgctgctcctgcagcgcacctctgcagccaggcg
tgtgtatgcctccatcctcaatgtgcgcaccaatacggatggtttcaaggagcagggcatcacataccctattggcaagatgcaaaatcgcctg
atccgcgagacctacgaggagattggtcttaaccccgccgatgtggtttacgtggaggcacacggtaccggaaccaaggtgggcgatcccc
aggaggtgaactctatcactgacttcttctgcaaggaccgtacgacccctctgttgatcggatcggtcaagtccaacatgggtcactcggaacc
cgcttccggtgtctgctctgtggccaagattctgatcgccatggaggagggcgtcattcccggtaacttgcactacaacaagccgaacccaga
tctttatggactcgttgatggacgtctcaaggtggtcgacaggaatctgccttggaacggtggcatcatcggcctgaactcctttggttttggagg
agccaacgctcacgttatcttaaagtcgaaccccaagcccaaggctttgacccccaaggatggagcactgaaagtggtcctcgcctctggac
gcacctttgaggctgtggagcagttgctcgagtccgcctctactaatgccgatgatgatgagtacctgcaactgatcaacgagatccactcaaa
ggctattccgaatcacttcttccgtggctacggagtggtgagcagcaagggcacccaccaacgtgaggtgatcgagtccaacgacgataaac
gccctatttggtacatctactccggcatgggcagccagtgggctagcatggccaaggatctgatgaagatcgaggccttcgccaagaccattc
agaggtgtgccgatgtcctaaagcccgagggagtggatctaatcgatgtcctgacccgttccacggacaagagcttcgaaaacattctgaact
ctttcatctcgattgctgccatgcaagtggctcttaccgatttgctcagctcgctgggcatccaccccgacggaattgtgggacactcggtggga
gagcttggatgcgcctacgccgatggctgcttcaccccggagcagaccgtgctggctgcttactggcgaggaaagagcattctggacactca
actggccaagggcaagatggccgctgtgggattgagctgggaagatgcgcacagccgtgtgcccagcgattgcttcccggtgtgccacaa
cagcgaggacaactgcaccatctctggcccggaggcttccatcgaggctctggtcgctaaactgaacgcggagggagtgttcgccaaggc
agtgaactcatccggctacgctttccacagcaagtacatcgctgaagctggacccaaactacgcaagagtctggagaagatcattcccaacg
ctaagaaccgtaccgcccgctggatcagtaccagcatcccagaatccgcttggaacacaccggtggccaagcaatcctcggccgcctacca
cgtgaacaatctgctgtcgccagtgctcttccacgaggcccttcagcacgttcccaagaacgccatttccgtggagattgctccccacggcctg
ctccaggctatcctcaagcgcgccctgggtcctgatgccaccaatctgagcttggtgaagcgcggacacgagaacaacgttgagttcttcctc
accaatgtgggcaaactctttgccgccggagcccagcctcaggtacttactctagtgcgtcctatcagctacccggtgggtcgtggaacccccc
atgttgaactccaaggtcggatgggaccacacccagaagtggctggtggccaagttcggcaaggaaacttcttctggtgagaccatcgtgga
agtcgatctctccaaggaggatgatgctttcctggctggacacacgatcgacggacgtatcctcttcccggccactggctacatgactctggct
tggcagacattcgctaagatgcagggaagtgagttccacaagaccccggtggtaatggagaacctcgtgttccaccgtgccactattcttaac
aagaacgcagtggtcaagtttggcattaacttcttcgatggaactggcgcttttgagatttgtgagagcggtagcttggcggtttctggaaaaata
accattccagagtctattgacaacgaggagttgcccctggaggagcaaactcccagcgctgttgccaaggagctaggcaccaacgatgtcta
caaggagctgcgcctccgaggctacgactatggtggaattttccgcggcattgttcgttctgacaccgtggcctccactggaaagcttcagtgg
gtggacaactggattagtttcatggacaccatgttgcagttcagtatccttagcaagaaccttcgtgagctgtacctgcccactcgcatcgagcg

agcggtgattaatcccgccaagcactttgaattgctgagcgccctcaccaaggaggaacaggttgagactggactacctgtgcagtggtaca
gcgacattaacgtgatcaagagtgcgggcgtggagctgcgcggtctaaaggccaatctcgcccagcgtcgtcctggcacacaggcacctcc
cactctggagcgctaccagtttgtgcccaacatcaacaccacggatctaaatgaaaactctgagaaggctcgcctgcacgccttggatgtggc
catccaggtgatcatcgagaactccagcggagccgtcaagctaaagggcgtggaattggccaatggacgcaaccccgacgttttggttgcc
aaccgcctgctgcagattatcgagggtgagcccgtgcttactggtgatgtggctgtggtcacctcgaacaacaatgaagagaccataaccgc
cgccctaggagattcaggagttcgagtggtgtccaaggacgtgctgaaggaaccggtagagcagaactgtcacttcgtctttggcatcgatgt
cctttcccgaccggacaccaagaccctggagaactccattgccagcattcgcgagaacggtttcctgatccttgaggagacgctgcccactta
caccaagacaggccgtgccctgctgaccaaatttggattcgttgccgttcaggaacagagtctgggagccacccgtgtcctggtcctcgccc
gcaaggctgttgatctgaaaactcgcaagtccgttgtggttgtggccaccgagcagaactttaactgggtggatgacttgaaggccgctctgg
ctactgctgctacggaggagcagtatgtgtacgtggtttgccagggagaggagctgtttggagctgtgggtctgatgacgtgcatcaagaacg
agaatggcggcaaactggctcgcctggtcttcgtgcaagatgccaaggctgagaagttctcccttacatcgaccctgtaccgccagcagctg
gagaaggacctcatctcgaacgtgctgaagaacggcgcttggggcactttccgtcacctgaaactggagacccagcaggccactctccaag
tggagcatgcctatgtaaacgctctggtcaagggtgatcttgcttccctcaagtggatcgaggctgcccaggcggataccgctgcaactgtcg
acaagaacctggaaacttgcaccgtctactatgcgcccatcaacttccgtgacgttatgttgacctctggtaagctggccgcggatgccttgcct
ggagatctggctgagcaagattgtgtgctgggtctggagttcgctggtcgcgatacccagggtcgtcgtgtaatggccatggtgccggctaaa
tcacttgctaccacctgcgtggcaagcaagcgcatgatgtggcaaatccctgagaagtggaccatggaagaggcttcgaccgttccttgcgtg
tactccaccgtctactacgctctggttgtgcgaggacaaatgaagaagggtgagaagatcctcattcacgctggctccggaggagttggtcag
gcggccatctctgtggctctggcccatggcttgaccgtgttcaccacggtgggcagcaaggaaaagcgcgagttcctgctgaagcgattccc
caaactgcaggagcgtaacatcggcaactcccgagacacctccttcgagcagttggtcctgcgcgagaccaagggacgcggagtcgatct
ggtgctgaactcgttatctgaagagaaactgcaggcctcgatccgctgtttgggtctgaatggacgcttcctggagattggcaagttcgatttga
gcaacaacagcccacttggaatgtccgttttcctcaagaacacctccttccacggcattttgcttgacagtgtgatggagggcgaggaggagat
gcaaaaccaggttgtctccctggtcgctgagggcatcaagaccggagccgttgtacccttccgacctcggtgtttaacgaccagcaggtgg
agcaggccttccgtttcatggcctctggcaagcacatcggcaaggtagtgatcaaggtgcgtgatgaggaggcaggcaagaaggcgctgc
agcccaagccgcgcctgatcaacgccattccacgcacatacatgcacccggagaagagttacatcctggttggaggtcttggcggattcggt
ttggagctgaccaactggttggtgacgcgtggagcccgctacattgtgctgacctctcgttccggagttaagaccggctaccagggtctgatg
atccgtcgctggcaggaacgtggcgtaaaggtggtgattgacaccagtgatgtgaccaccgccgctggagctaaaaagctgctggagaata
gtaacaagctggctttggtcggaggcatcttcaacctggcggctgttctccgcgatgctctgatcgaggatcagaccgccaaggacttcaaga
cagtggctgatcccaaggtgacggccaccaagtacctggatcagttctctcgcgatatctgcactgagctggactacttcatctgcttctccagt
gtttcctgcggtcgtggtaacattggtcagaccaactacggattggccaactctgcaatggagcgaatttgcgagcagcgccaggtgagcgg
attcccaggcaccgccatccaatggggagccatcggagacaccggtctggtgctggagaacttgggtgacaatgacaccgtgattggagga
accctgccccagaggatgccctcttgcctgcagaccatcgacctgttcctgcaacagccccatccggttgttgcctccatggttgtggccgag
aagcgcaagtcggatcagtcggctggagtcagcctaattgccaccattgccaacatccttggtctccgggacaccaagaatatccaggacgg
agcatcgcttgctgatctcggaatggactccctgatgagcgctgagatcaaacagactttggagcgaaactttgacattgttctgtccgcccag

gagatccgtcagctcacctttggagccctcaaggccatggatggtggtgctgatgtcaagccagccgcagctgctcccgcagccgctgctgg
agtcccagaggcaaacattacctctggtggatcctcgcgcacagctagtcccatgggcgatggaacgcaggtggtgttcaccacgtccctca
tccccaccgaggcaattgtacaactggacacaaaggcacctgcaaacagcaaacagagccccatcttcttcatttccccgatcgaggggttcg
cctccgcactggagccgctggccaagcgattggaggtgcccgcctatggactgcagtacacggaagccgtgccctccgattctctggaatc
agcagccaaattcttcatcaagcagctgcgcaccgtccagccaaagggtccctacaaactagccggatactccttcggttgcctgctcaccta
cgtgatggccggtattctggaagagaccaacgaggtggccaatgtgatcatgctggacggagccccaagctacgtcaactggtacacgagc
agcttcaagcagcgttacactgacggcaccaacgccgataacgacaaccagagctacggcttggcctactttggaatcgtgttggccaacatt
gactacaaagcgctggtgcgtctgctaatagtgatccccacatgggaggagaaactcgagaggttcgctgagctgatgagcaatgagatcac
ccagcccgtggaaacgatcaagaaatccgccactctgttctacaagaaactggagctggccgatggctaccagcccactctgaagctcaaga
cgaatgtgaccttggtcaagcccacagacaactctgccaagctggacgaggactatcgccttaaggaggtctgcacaaagcccgtggaagt
acacactgttgagggcaaccaccgcaccttcctcatcgaggatcagtccctgaagaccatccagagcatactgaagcgtctgttcaactaa

>DmPPT

Atgctgccgaaatacagttgtgtgcgctgggcgttagatttatcacagtggcgcccgacacttgccgagctcagcttagcgactgcatgtttac
aagccgaggagaaggctcgtctcatgaaattctacttccgggatgatttcgatgcatccttaatagggcgcctgctcatgcggaaatttgtgcg
ggacacaaccggaatcgcttacgatcaaataagattcgagcgtgatggacgcggcaaacccttcctggtcaatagcgtcttgggaccgtgtc
ggatcgacttcaacgtttcccatcagggcaactatagcgtgctggccggaatcgttgagagatcaattgcagacgagccacctgtgaaaatcg
gaatcgatgttatgaaaattgagtattcaggtggaaagccattaaccgaattcttccgcctgatgacgaggaacttttccgacgccgagtggag
atttatccggagccgacagtctgaggccgagcaattaaaagcgtttatgcgattctggtgtctgaaggagagttacgtcaaaaatgtcggtgttg
ggattaccatgaatttacagaaaataagtttcaaggtgaagactgacgacgtagcgaaagggaaggtggttcaggacacggtcttggaggtc
gatgggaatttactgacgaattggtgttttgaggagcatctgctggacgagcagcacagcgtagccatagcatttgaaaatttaggaaaaatgg
cagatgaaggcacatttaaagttttaggatatgaagagttgatggaaggcgcattgcccctaatggaaaaagataattcttattgtcaaagggttt
tggaaaagcaatataaacgtttccaataa

>SFP

Atgaagatttacggaatttatatggaccgcccgctttcacaggaagaaaatgaacggttcatgactttcatatcacctgaaaaacgggagaaat
gccggagattttatcataaagaagatgctcaccgcaccctgctgggagatgtgctcgttcgctcagtcataagcaggcagtatcagttggacaa
atccgatatccgctttagcacgcaggaatacgggaagccgtgcatccctgatcttcccgacgctcatttcaacatttctcactccggccgctgg
gtcattggtgcgtttgattcacagccgatcggcatagatatcgaaaaaacgaaaccgatcagccttgagatcgccaagcgcttcttttcaaaaac
agagtacagcgaccttttagcaaaagacaaggacgagcagacagactatttttatcatctatggtcaatgaaagaaagctttatcaaacaggaa
ggcaaaggcttatcgcttccgcttgattcctttcagtgcgcctgcatcaggacggacaagtatccattgagcttccggacagccattccccatg
ctatatcaaaacgtatgaggtcgatcccggctacaaaatggctgtatgcgccgcacaccctgatttccccgaggatatcacaatggtctcgtac
gaagagcttttataa
